# Gene expression patterns of the developing human face at single cell resolution reveal cell type contributions to normal facial variation and disease risk

**DOI:** 10.1101/2025.01.18.633396

**Authors:** Nagham Khouri-Farah, Emma Wentworth Winchester, Brian M. Schilder, Kelsey Robinson, Sarah W. Curtis, Nathan G. Skene, Elizabeth J. Leslie-Clarkson, Justin Cotney

## Abstract

Craniofacial development gives rise to the complex structures of the face and involves the interplay of diverse cell types. Despite its importance, our understanding of human-specific craniofacial developmental mechanisms and their genetic underpinnings remains limited. Here, we present a comprehensive single-nucleus RNA sequencing (snRNA-seq) atlas of human craniofacial development from craniofacial tissues of 24 embryos that span six key time points during the embryonic period (4–8 post-conception weeks). This resource resolves the transcriptional dynamics of seven major cell types and uncovers distinct major cell types, including muscle progenitors and cranial neural crest cells (CNCCs), as well as dozens of subtypes of ectoderm and mesenchyme. Comparative analyses reveal substantial conservation of major cell types, alongside human biased differences in gene expression programs. CNCCs, which play a crucial role in craniofacial morphogenesis, exhibit the lowest marker gene conservation, underscoring their evolutionary plasticity. Spatial transcriptomics further localizes cell populations, providing a detailed view of their developmental roles and anatomical context. We also link these developmental processes to genetic variation, identifying cell type-specific enrichments for common variants associated with facial morphology and rare variants linked to orofacial clefts. Intriguingly, Neanderthal-introgressed sequences are enriched near genes with biased expression in cartilage and specialized ectodermal subtypes, suggesting their contribution to modern human craniofacial features. This atlas offers unprecedented insights into the cellular and genetic mechanisms shaping the human face, highlighting conserved and distinctly human aspects of craniofacial biology. Our findings illuminate the developmental origins of craniofacial disorders, the genetic basis of facial variation, and the evolutionary legacy of ancient hominins. This work provides a foundational resource for exploring craniofacial biology, with implications for developmental genetics, evolutionary biology, and clinical research into congenital anomalies.

## Main

Craniofacial development orchestrates the formation of the human face through the interplay of multiple cell lineages. These cell types, including mesenchyme, ectoderm, endothelium, and cranial neural crest cells (CNCCs), differentiate into a diverse array of tissues such as bone, cartilage, muscle, skin, and vasculature^1–3^. Together, these cells and tissues give rise to the face’s essential functions like respiration, mastication, communication, and sensory perception^4–7^, Disruptions to craniofacial developmental processes rank amongst the most common causes of human congenital anomalies, with orofacial clefts representing a significant portion of global birth defects^8–11^. Thus, there exists significant need to understand the molecular, genetic, and cellular mechanisms underlying craniofacial development in humans.

Studies utilizing model organisms, particularly mice, have offered key insights into craniofacial development and abnormalities^12–16^. However, significant differences exist between mouse and human craniofacial development, including variations in timing, cellular contributions, and gene regulatory networks^1,13,17–22^. Furthermore, human craniofacial features exhibit evolutionary adaptations that distinguish them from other mammals and primates, underscoring the necessity for human-specific studies ^5,13,21,22^.

Advances in single-cell and single-nucleus RNA sequencing (scRNA-seq and snRNA-seq respectively) technologies have enabled detailed characterization of cellular diversity and gene expression during development^12,15,23–27^. These tools are particularly valuable for resolving the dynamics of rare or transient cell populations, such as CNCCs, that play critical roles in craniofacial formation^13^. While previous efforts have developed single-cell atlases for murine craniofacial tissues, corresponding human datasets have been limited by sample availability, insufficient temporal resolution, and challenges in profiling craniofacial-specific populations^12,15,23^. Only a few studies have examined bulk gene expression patterns and regulation specifically during human craniofacial development^13,14,28–31^, and only two datasets are currently available during the embryonic period of human development^13^. While mapping of human genetics findings to mouse craniofacial cell types has indicated potential disease-causing subtypes^32^, the limited number of replicates underlying the mouse data and differences between human and mouse craniofacial development preclude confident interpretation.

To address these gaps in knowledge, we constructed a time-resolved gene expression atlas of human craniofacial development when the bulk of human craniofacial development occurs^33,34^. Using snRNA-seq on craniofacial tissues from 24 individual human embryos encompassing six key time points from 4 to 8 post-conception weeks, we profiled over 42,000 nuclei and identified seven major cell types, including mesenchyme, ectoderm, endothelium, muscle progenitors, and CNCCs. Integration with human spatial transcriptomics further validated the localization of these subtypes within the developing human face. Comparative analysis with murine craniofacial datasets generated here and previously published^23^ highlighted significant conservation of major cell types and their gene expression programs, alongside species biased markers that reflect differences in mouse and human biology.

Beyond the developmental biology of craniofacial formation, this study explores the genetic and evolutionary factors shaping human craniofacial features. By integrating genome-wide association studies (GWAS) with our atlas, we identified cell type-specific enrichments for genetic variants associated with normal facial variation. We found that specific subtypes of ectoderm and mesenchyme, likely spatially restricted, contribute to different aspects of facial appearance and shape. We also examined rare variants associated with congenital craniofacial disorders such as orofacial clefts. We find that *de novo* protein damaging variants identified in orofacial clefting trios are enriched in genes that specify distinct cell subtypes in the face. This enrichment was heavily biased toward ectodermal subtypes that is largely obscured in previous analyses based on bulk chromatin and gene expression^13,35–38^. We find the damaging variants coalesce in the ectodermal derived nasal placode implicating this early structure in orofacial clefts. Our analysis also uncovered evidence linking Neanderthal-introgressed sequences to genes with biased expression in specific craniofacial cell types.

This comprehensive atlas provides a high-resolution view of the cellular and molecular landscape of human craniofacial development, integrating gene expression, spatial mapping, and evolutionary genomics. Our work not only enhances our understanding of human craniofacial biology but also establishes a framework for future studies aimed at uncovering therapeutic targets and evolutionary insights into one of the most defining features of human anatomy. This data can be explored through an interactive web application that is accessible to most researchers: https://cotneyshiny.research.chop.edu/shiny-apps/craniofacial_all_snRNA/ as well as alongside the growing number of single cell datasets hosted at the Chan-Zuckerberg CellXGene Discover resource^39^.

## Results

### Time-resolved atlas of gene-expression in the developing human face

To characterize the cellular landscape of human craniofacial development we performed snRNA-seq analysis of 24 individual human embryos across 6 distinct time points, encompassing major milestones of human craniofacial development from 4 to 8 post conception weeks (Fig. 1A). We profiled the entire craniofacial prominence from multiple biological replicates at each time point resulting in 86,359 individual nuclei after filtering for doubles and quality of per nucleus data. While experiments in mouse offer precise control of tissue sampling for downstream processing, samples obtained from human embryos are more difficult to control what is collected. To identify potential biases or nuclei obtained from extraneous tissues we performed initial clustering of all samples to identify potential extraneous cell types. This analysis revealed a total of 13 distinct clusters (Fig. S1a). This number of clusters was substantially higher than the main cell types identified in mouse craniofacial developmental studies, suggesting that the human samples potentially contained extraneous tissues that are not part of the craniofacial complex^12–14^. When we examined the contributions of individual samples to these clusters, we found several clusters that were made up of nuclei derived from a small number of samples (Fig. S1b). Closer inspection of the genes strongly expressed in these clusters revealed many canonical neuronal genes, such as *TUBB3* and *MAP2* (Fig. S1c-d). We reasoned these clusters were derived from developing brain tissue not directly part of the craniofacial structures. We therefore excluded these nuclei from downstream analyses resulting in a total of 42,131 remaining nuclei with an average of 4095 nuclei from each sample (Fig. S2a) and a median of 7500 counts from 2250 genes per nucleus (Fig. S2b and c). We observed that early samples had consistently higher mitochondrial reads (Fig. S2d), potentially reflecting their higher dependence on mitochondrial output or an artifact related to lower cell numbers in the processing of each sample.

**Figure 1.**
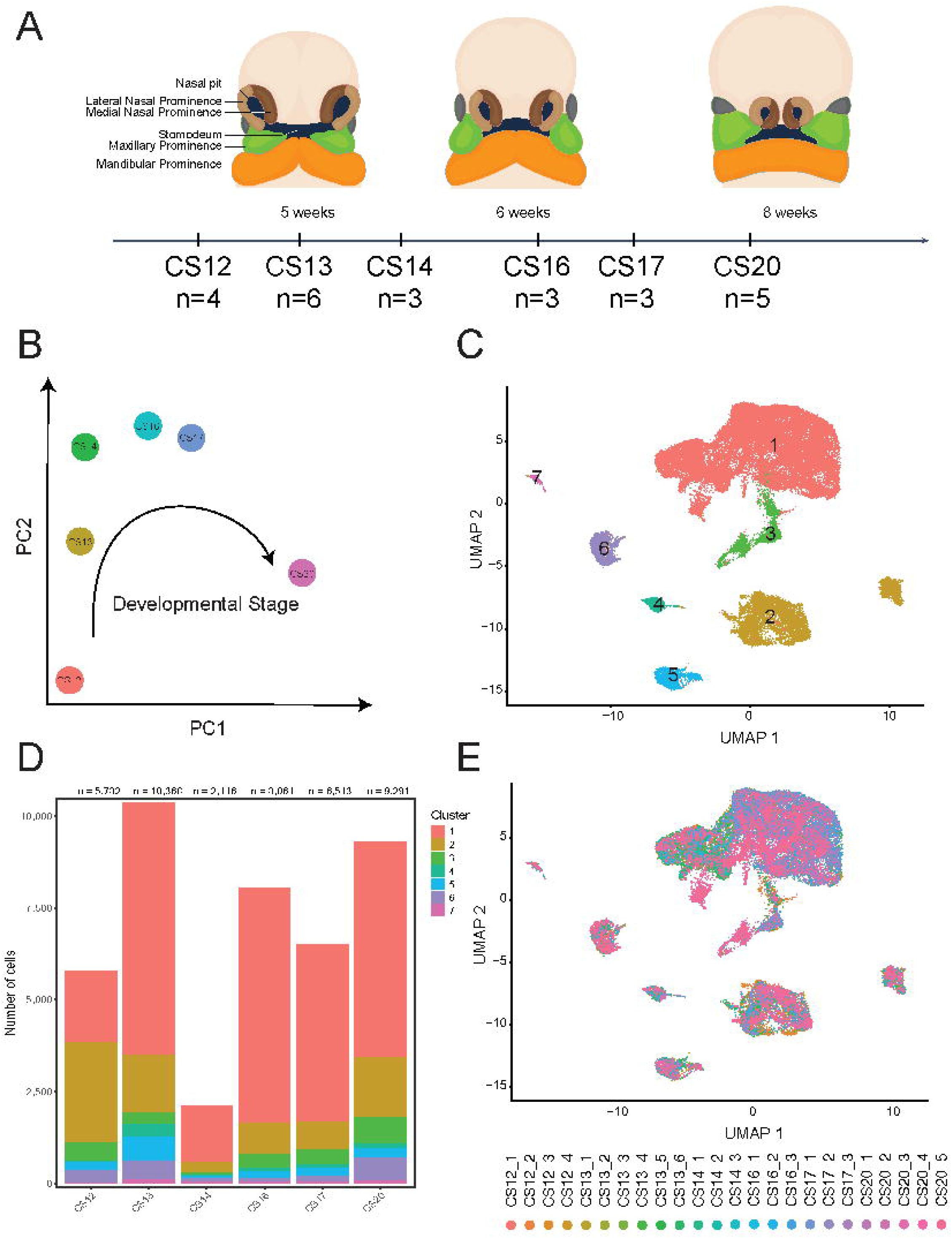
Generation of single nucleus gene expression atlas of human craniofacial development. A). Anatomical regions of the developing craniofacial region from 4 to 8 weeks post conception. Individual Carnegie Stages (CS) and replicates at stage are indicated below images. B). Pseudo-bulk gene expression of tissues from each stage displayed in principal component (PC) space based on the first two PCs. Progression of developmental time is indicated along PC1 dimension. C) UMAP projection and cluster identification of all human craniofacial cells after filtering of neurons. D). Number of cells obtained at each CS stage for each cluster identified in C. E). Distribution of samples from each sample across the UMAP projection.

To determine the quality of these filtered samples we sought to compare to other well characterized gene-expression profiles of craniofacial development. Our previous studies of bulk gene expression during human craniofacial development revealed a strong time related component across the samples^13^. When we combined gene expression profiles from all nuclei of a specific stage into pseudo-bulk gene expression profiles individual replicates were well correlated with others at the same time-point and less so with samples with greatest differences developmental time across this time course (Fig. S3). Principal component analysis of these pseudo-bulk profiles largely recapitulated our previous results with the first principal component ordering samples readily by known stage of development (Fig. 1B, Fig. S4a). Furthermore, when we performed differential expression between the pseudobulk samples we found very similar results to those obtained by bulk gene expression between the same timepoint comparisons (Fig. S4b). Specifically, the greatest number of differentially expressed genes were observed between the earliest and latest timepoints that could be compared across the two data sets (CS13 vs CS17) (Fig. S4c). Overall, these results suggest that our single-nucleus expression data closely resembles the bulk gene expression data that we have previously shown is enriched for many aspects of craniofacial biology and developmental abnormalities relative to many other tissues and cell types^13^.

### Identification of major cell types present in craniofacial development

Having established that the single nuclei profiles at the pseudobulk level captured many of the expected aspects of craniofacial biology, we proceeded to re-cluster the filtered nuclei to first identify the major cell types present in the developing face. We identified seven major clusters and projected these high-dimensional data into two dimensions using Uniform Manifold Approximation and Projection (UMAP)^40^ (Fig. 1C). The clusters were contributed to by samples from each of the replicates and stages in very similar proportions (Fig. 1D and E). Interestingly, this was two more distinct clusters than previously characterized in the E11.5 mouse craniofacial structures^12^. We reasoned that this could be due to differences in human and mouse development, but most likely related to the additional replicates and timepoints and how tissues were collected and processed. To address this, we first examined expression of the five genes examined by Li et al, *ALX1*, *EPCAM*, *HEMGN*, *CDH5*, and *FCERG1* as markers of mesenchyme, ectoderm, blood, endothelium, and immune cells respectively (Fig. 2A). *ALX1* was most strongly expressed in cluster 1, *EPCAM* in cluster 2, *HEGMN* in cluster 4, *CDH5* in cluster 5, and *FCERG1* in cluster 7, while clusters 3 and 6 did not show signal for any of these genes. While some of the timepoints were derived exclusively from female and male samples, CS12 and CS13 respectively, we did not observe any significant bias is these main cluster (Figure S2E).

**Figure 2.**
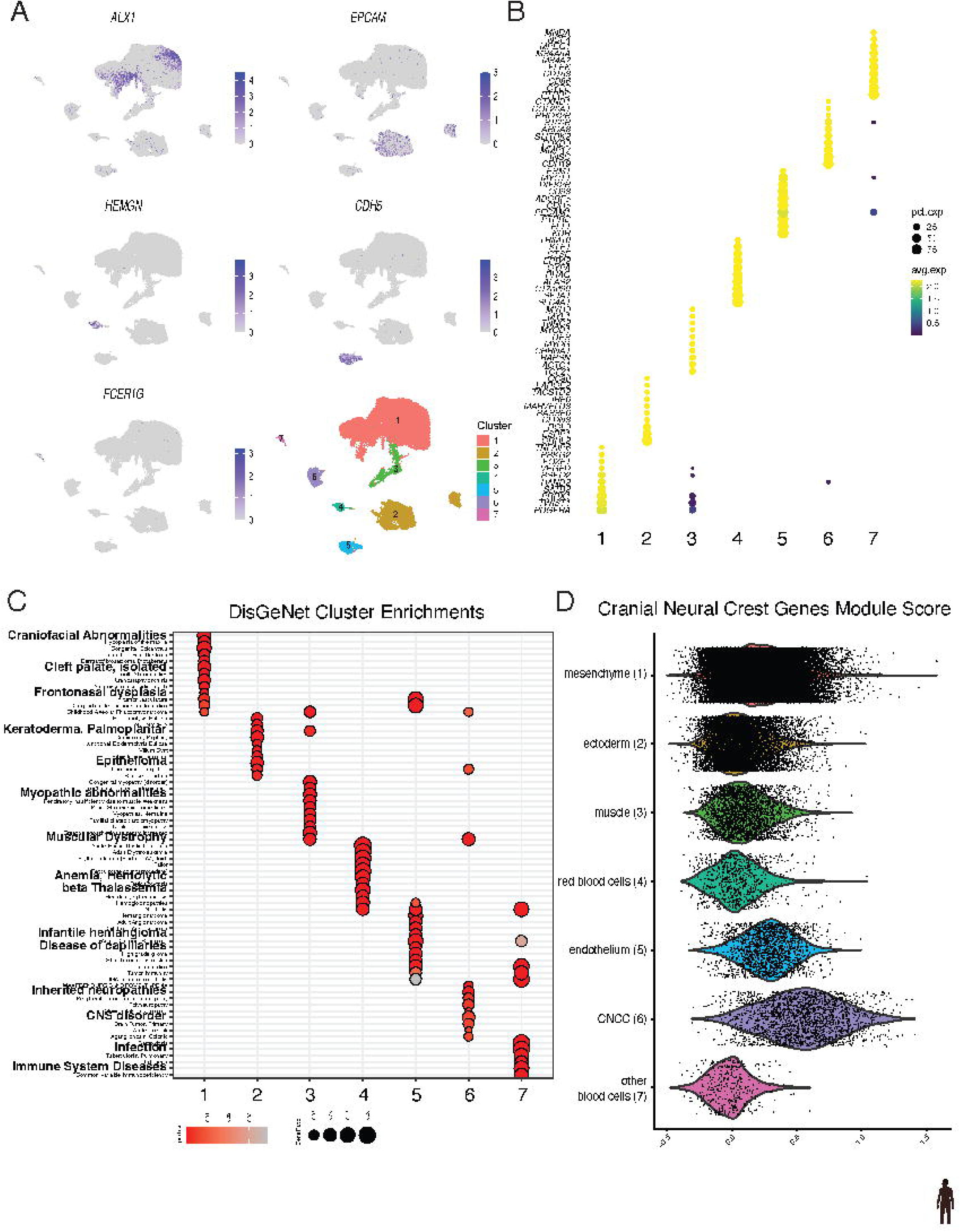
Identification of main cell types in the developing human face. A). Gene expression feature plots for indicated genes across UMAP projection. B) Average and percent expression for the top 10 maker genes for each main cluster. C). Disease ontology enrichments of categories curated by DisGeNet for each indicated cluster. D). Identification of CNCC cluster based on module score of curated neural crest genes and labelling of all remaining clusters. Violin plots and individual values for all cells of a given cluster type based on CNCC module score calculated by Seurat.

In an attempt to characterize these unknown clusters we first identified the top 10 genes that were most strongly differentially expressed between the individual clusters (Fig. 2B, Supplemental Table 1). Cluster 1 was marked by *PDGFRA, TWIST1, and PRRX1,* consistent with identifying this cluster as mesenchyme^41,42^. Cluster 2 was identified by *GRHL2* and *ESRP1*, genes that have been reported to be specifically active in surface ectoderm and epithelial cells^43–46^. Cluster 4 was marked by *SPTA1, ALAS2, RHAG,* genes involved in erythrocyte function^47–51^. Cluster 5 showed highly biased expression for *KDR and FLT1,* genes associated with the vascular system and endothelium function^52,53^. Cluster 7 was marked by *PTPRC*, *CD86,* and *CD136,* consistent with immune cell function^54–56^. These all confirmed the initial identities suggested by the markers described by Li et al in E11.5 mouse craniofacial tissue. The unknown cluster 3 showed highly biased expression of *MYOG*, *MYL1*, and *MYH3*, all genes related to muscle specification and function^57–59^. The unknown cluster 6 showed strongly biased expression for *CDH19*, *INSC*, and *MMP17*. These genes are involved in a variety of biological processes including cell adhesion, spindle orientation during mitosis, and degradation of extracellular matrix^60–62^. We also noted specific expression of *FOXD3,* a developmental transcription factor which has been linked to pluripotency maintenance in stem cells and specification of neural crest in multiple species^63–66^.

We then analyzed the top 100 marker genes from each cluster for gene ontology, pathway, and disease enrichments. The genes that identified putative mesenchyme cluster 1 were enriched for a number of biological process categories related to skeletal, cartilage, and roof of mouth development (Fig. S5A, Supplemental Table 2); cellular components related to collagen processing (Fig. S5B, Supplemental Table 3); molecular functions related to gene expression, extracellular matrix, and collagen binding (Fig. S5C, Supplemental Table 4); pathways related to production of extracellular matrix (Fig. S5D, Supplemental Table 5); and diseases including cleft palate and frontonasal dysplasia (Fig. 2C, Supplemental Table 6).

Genes most strongly expressed in cluster 2, likely ectoderm, were enriched for biological process categories related to tight junction assembly and cell adhesion (Fig. S5A, Supplemental Table 2); cellular components related to the plasma membrane (Fig. S5B, Supplemental Table 3); molecular functions related to cadherin and laminin binding (Fig. S5C, Supplemental Table 4); pathways related to tight junction and Hippo signaling (Fig. S5D, Supplemental Table 5); and diseases including epithelioma and keratoderma (Fig. 2C, Supplemental Table 6).

Putative muscle progenitor cluster 3 marker genes were enriched for biological processes related to muscle cell differentiation (Fig. S5A, Supplemental Table 2); cellular components of the sarcomere (Fig. S5B, Supplemental Table 3); molecular functions related to actin filament binding (Fig. S5C, Supplemental Table 4); calcium signaling pathways (Fig. S5D, Supplemental Table 5); and diseases including myopathic abnormalities and muscular dystrophy (Fig. 2C, Supplemental Table 6).

The markers of red blood cell cluster 4 were enriched for biological processes related to erythrocyte homeostasis and oxygen transport (Fig. S5A, Supplemental Table 2); cellular components of the hemoglobin complex (Fig. S5B, Supplemental Table 3); molecular functions related to heme and oxygen binding (Fig. S5C, Supplemental Table 4); pathways involved in Malaria response and mineral absorption (Fig. S5D, Supplemental Table 5); and diseases including hemolytic anemia and beta thalassemia (Fig. 2C, Supplemental Table 6).

Genes with highest expression in putative endothelium cluster 5 were enriched for biological processes related to endothelial cell differentiation and proliferation (Fig. S5A, Supplemental Table 2); cellular components including plasma membrane rafts and caveola (Fig. S5B, Supplemental Table 3); molecular functions related to Notch and guanyl nucleotide binding (Fig. S5C; Supplemental Table 4); pathways involved in fluid shear stress and atherosclerosis (Fig. S5D; Supplemental Table 5); and diseases of the capillaries and hemangiomas (Fig. 2C, Supplemental Table 6).

Immune related cluster 7 marker genes were enriched for biological processes related to cytokine production and immune response (Fig. S5A, Supplemental Table 2); cellular components including specific and tertiary granule membranes (Fig. S5B, Supplemental Table 3); molecular functions related to Toll−like receptor binding and immunoglobulin receptor activity (Fig. S5C, Supplemental Table 4); pathways related to the phagosome and complement and coagulation cascades (Fig. S5D, Supplemental Table 5); and diseases including many types of infections and immunodeficiencies (Fig. 2C, Supplemental Table 6).

We then turned to the not yet concretely identified cluster 6. The marker genes we identified for this cluster were enriched for biological processes related to glial cell differentiation and myelination (Fig. S5A, Supplemental Table 2); cellular components including plasma membrane signaling receptor complexes and exocytic vesicles (Fig. S5B, Supplemental Table 3); molecular functions related to protein tyrosine kinase activator activity (Fig. S5C, Supplemental Table 4); and diseases related to central nervous system disorders and neuropathies (Fig. 2C, Supplemental Table 6). We did not detect any significant pathway enrichments for this particular cluster.

### Identification of presumptive human cranial neural crest

The marker gene ontology analysis successfully confirmed the identity of six of the seven major clusters. However, cluster 6 remained difficult to identify due to the variety of enrichments identified amongst marker genes. Beyond the more nervous system-oriented enrichments listed above we also found significant biological and disease enrichments that were shared with the mesenchyme and muscle clusters. This included enrichments for extracellular matrix organization and binding, cell adhesion via plasma-membrane, skeletal muscle system development, and several types of tumors (Fig. 2C, Fig. S5A-D, Supplemental Tables 2-6). When we more closely inspected biological process categories identified for cluster 6, we observed additional enrichments related to Schwann cell development and melanocyte differentiation (Supplemental Table 2). Closer inspection of full disease enrichments for this cluster revealed several types of Waardenburg Syndrome, Hirschsprung Disease, and demyelination disorders (Supplemental Table 6). The wide variety of biological functions and specific disease enrichments all suggested that this cluster might be enriched for neural crest cells. Marker genes driving these enrichments included *EDNRB*, *ERBB3*, *PAX3*, *SOX10*, *SPP1*, *TFAP2B*, and *ZEB2*, genes well known to be involved in various aspects of neural crest specification and migration^67^. However, while these genes are biased toward cluster 6 relative to other clusters, they are not exclusively expressed in cells found in this cluster (Fig. S6A). Amongst these *ZEB2* is more broadly expressed across all clusters except for ectoderm. Further inspection of marker genes revealed that while some of these genes are indeed strongly biased toward cluster 6, only a small percentage of cells from this cluster express each gene (Fig. S6B). Qualitatively, expression of each of these genes could be observed outside of cluster 6 and potentially in subclusters of the main clusters we have defined thus far (Fig. S6C). Given the heterogeneity of expression of each of these marker genes we reasoned that jointly analyzing expression of a module of genes might be a better indicator of cell type identity as has been shown in other single cell-based studies^68^. When we examined a module of genes from regulatory networks recently identified in cultured human and chimpanzee cranial neural crest cells (CNCCs) ^69^, we found significantly higher expression in cluster 6 (Fig. 2D). Together these results strongly point to this cluster being enriched for putative CNCCs.

Thus far few studies have been able to identify significant populations of CNCCs from primary human tissue^70–72^. To better understand the gene expression programs that are active in these cells we first performed subclustering on these cells (n = 1821). We identified 11 distinct subclusters from this original population (Fig. 3A). The seven main clusters derived from the largest numbers of cells were annotated as CNCCs, while the four more punctate clusters were initially annotated as CNCC like (cnl). These clusters consisted differentially of cells derived from each of the stages profiled. Those clusters that were heavily biased toward CS12 were labeled as early (eCNCC), those that were biased toward CS13-C16 as intermediate (iCNCC), and those that were derived primarily from the CS17 and CS20 timepoints as late (lCNCC) (Fig. 3B). When we examined gene expression of many CNCC markers from the literature we found variable patterns of expression. *SOX10* expression was observed in all of the CNCC and cnl clusters along with *NR2F1* and *NR2F2,* two genes identified as master regulators of CNCC fate^6,73^ (Fig 3C). *TFAP2A* was expression was observed in all clusters but was considerably lower in the late CNCCs. Its ortholog *TFAP2B* was conspicuously absent from one late CNCC cluster (lCNCC2) and from cluster cnl2 (Fig. 3C). *ETS1* and *FOXD3* were generally expressed in most subtypes, but both expressed at very low levels in lCNCC1 (Fig. 3C). PAX3 expression was more variable but in populations distinct from the two previously mentioned transcription factors. *SOX9* and *COL20A1* were more specifically expressed across the clusters, but again in non-overlapping patterns.

**Figure 3.**
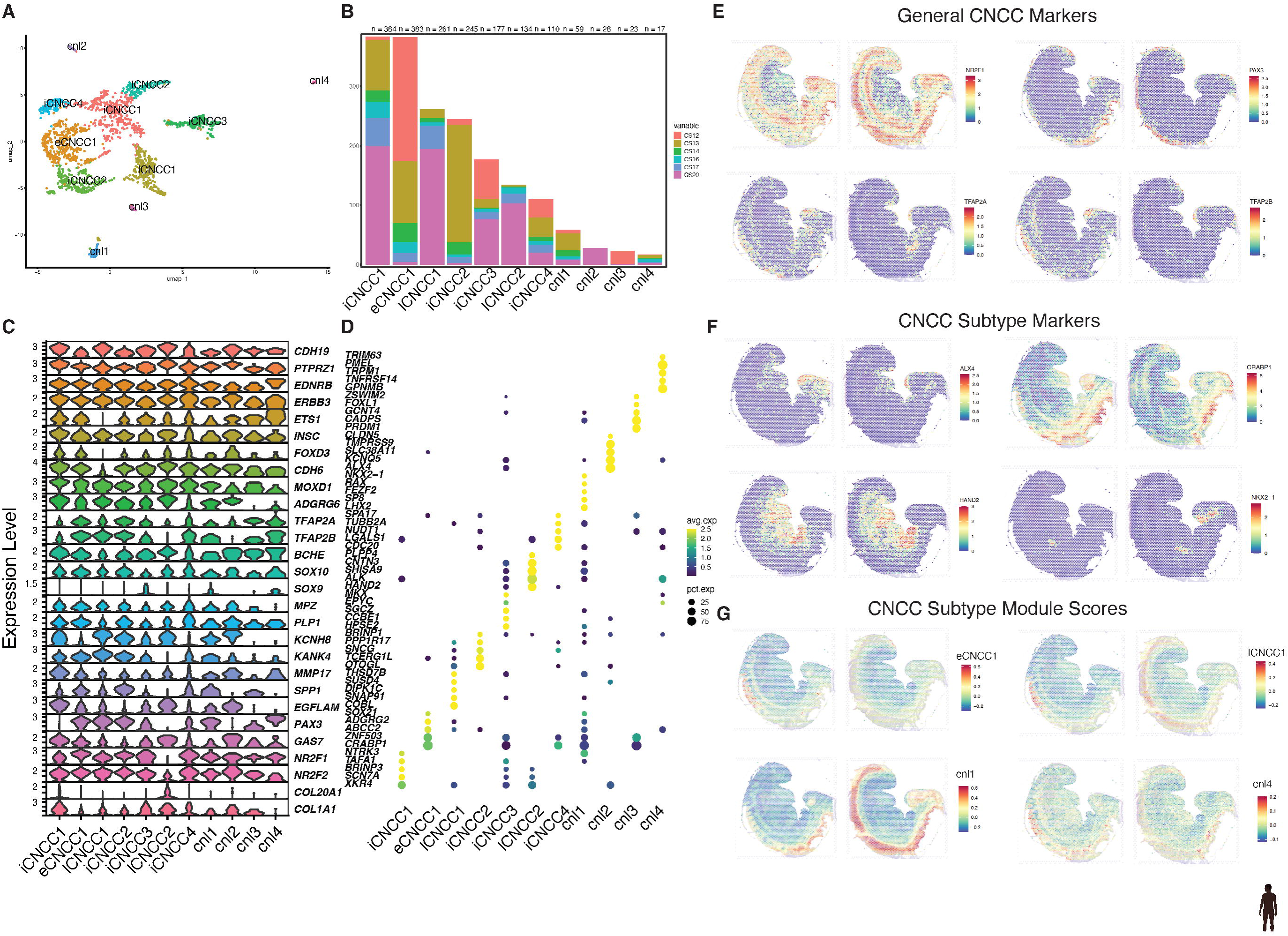
Identification of CNCC subtypes in the developing human face. A). UMAP projection of subclustered CNCC main cell type. B). Contribution of cells from each CS timepoint to each CNCC subcluster. C) Violin plots of published neural crest marker genes across each subcluster. D). Average and percent expression for the top 5 marker genes for each of the CNCC subclusters. E). Gene expression spatial feature plot for indicated CNCC marker genes in two sections from a CS13 human embryo. F). Gene expression spatial feature plot for indicated CNCC subtype marker genes in same sections as E. G). Spatial feature plot for modules scores calculated from top 100 marker genes from indicated CNCC subtype.

Overall, these genes largely confirmed that the cells we identified have neural crest character, however they did not display distinct patterns across the clusters precluding easy identification of these putative subtypes. To identify genes that readily identified each of these subtypes, we repeated the marker gene identification performed on the main types above. We identified approximately 2000 genes that were differentially expressed across these subclusters with an adjusted p-value cutoff less than 0.05 and a log_2_ fold change greater than one (Supplemental Table 7). The top five marker genes in each subtype revealed multiple transcription factors that distinguish each cluster. These included *SOX21* in early CNCCs, *MKX* in intermediate CNCCs, *HAND2* in late CNCCs, and *NKX2-1*, *ALX4*, and *FOXL1*, in clusters cnl1, cln2, and cnl3, respectively (Fig. 3D). Identification of enriched gene ontology categories for each subtype revealed distinct functions for each. Marker genes of early CNCCs were enriched for process related to early pattern specification and axon guidance. Intermediate CNCC clusters were enriched for functions related to extracellular matrix organization and skeletal system development. Late CNCC clusters were enriched for various channel activity and sympathetic nervous system development. The cnl1 cluster was enriched for several categories shared with eCNCC1 suggesting this was an early multipotent neural crest type. The cnl4 cluster was very specifically enriched for functions related to pigment granules and melanin biosynthesis indicating these were melanocytes, a cell-type derived from neural crest (Figure S7, Supplemental Tables 8-11).

Thus far our analysis has lacked spatial localization, making it unclear where these cell types are derived or reside in the intact human embryo. Recently published spatial transcriptomics on two sections of a human CS13 provided an opportunity to identify such patterns of expression^25^. We reprocessed this data, merged all the cells from both sections, identified cell types, and confirmed their spatial locations (Figure S8). We then examined expression of marker genes that identified CNCCs versus the other major craniofacial cell types. *NR2F1* was broadly expressed across the embryo whereas *PAX3*, *TFAP2A*, and *TFAP2B* were more regionally restricted to the head and neural tube regions (Fig. 3E). Genes identified as markers of CNCC subtypes showed a variety of patterns of expression. *ALX4* was generally restricted to the head region and putative frontal nasal process region. *CRABP1* was found in the anterior neural tube, eye region, and the limb. *HAND2* expression was observed in putative pharyngeal arch regions, heart, and limb. *NKX2-1* had highly restricted expression in a location that could represent a fusion zone between the lateral nasal prominence and the maxillary prominence (Fig. 3E). We then calculated module scores on these spatial data using the marker genes from each of the CNCC subtypes. We found that at this stage of development, each of these sets of marker genes were generally biased toward the neural tube region of the embryo with cnl1 marker genes showing the most restricted pattern of expression.

### Conservation of cell types and gene expression programs in human and mouse craniofacial development

The analysis above showed compelling evidence of the identities of the major cell types in the developing human face. This included two cell types, muscle and CNCCs, not previously observed in single cell atlases of mouse craniofacial development^12,15,23^. We wondered whether these cell types were not present in these mouse datasets due to sampling differences in tissues and broader timepoints. To address this, we generated single-nucleus gene expression data from mouse craniofacial tissues harvested from multiple biological replicates of E10.5 to E12.5. These samples reflected the major morphological landmarks of the human tissue profiled allowing a more direct comparison of cell types. We then further combined this data with recently published single cell gene expression data from E13.5 and E15.5 resulting in 79402 expression profiles after similar quality control filters applied to human data (Methods). When we clustered these data using approaches identical to the human data, we obtained the same number of main clusters with remarkably similar cluster ratios and organization in the UMAP projection space (Fig. 4A, Supplemental Table 12). When we examined gene expression of the same major markers profiled in human (Fig. 2A) we readily identified the same major mouse cell types including muscle and putative CNCCs (Fig. 4B). Roughly 70% of the tissue was of mesenchymal origin, 15% was ectodermal, and the remaining 15% was distributed similarly across the remaining 5 cell types. When we projected these cell types on our recent analysis of spatial gene expression in mouse E15.5 craniofacial sections, we found expected patterns of cell type localization (Fig. 4C). To determine if these cell types were specified by the same sets of genes, we compared marker gene identities obtained in the same fashion in both species. We found significant sharing of marker genes between the orthologous major cell types (Fig. 4D). The highest degree of sharing was found between mesenchyme, followed by endothelium and ectoderm. While still significant, the lowest degree of marker gene conservation was observed between CNCCs of each species (Fig. 4D). When we examined the functionally conserved marker genes for ontology enrichments, we observed distinct patterns of enrichment that confirmed the cell type assignments in each species (Fig. S9A-E, Supplemental tables 13-17). Disease enrichments related to general craniofacial abnormalities were observed in mesenchyme, while enrichment of cleft upper lip was observed in conserved markers of mesenchyme and ectoderm (Fig. S9F, Supplemental Table 18). These enrichments were driven by well-known craniofacial genes including *ALX1*, *ALX4*, *MSX1*, *RUNX2*, and *TWIST1* reinforcing their conserved role in mammalian craniofacial development (Supplemental Table 18).

**Figure 4.**
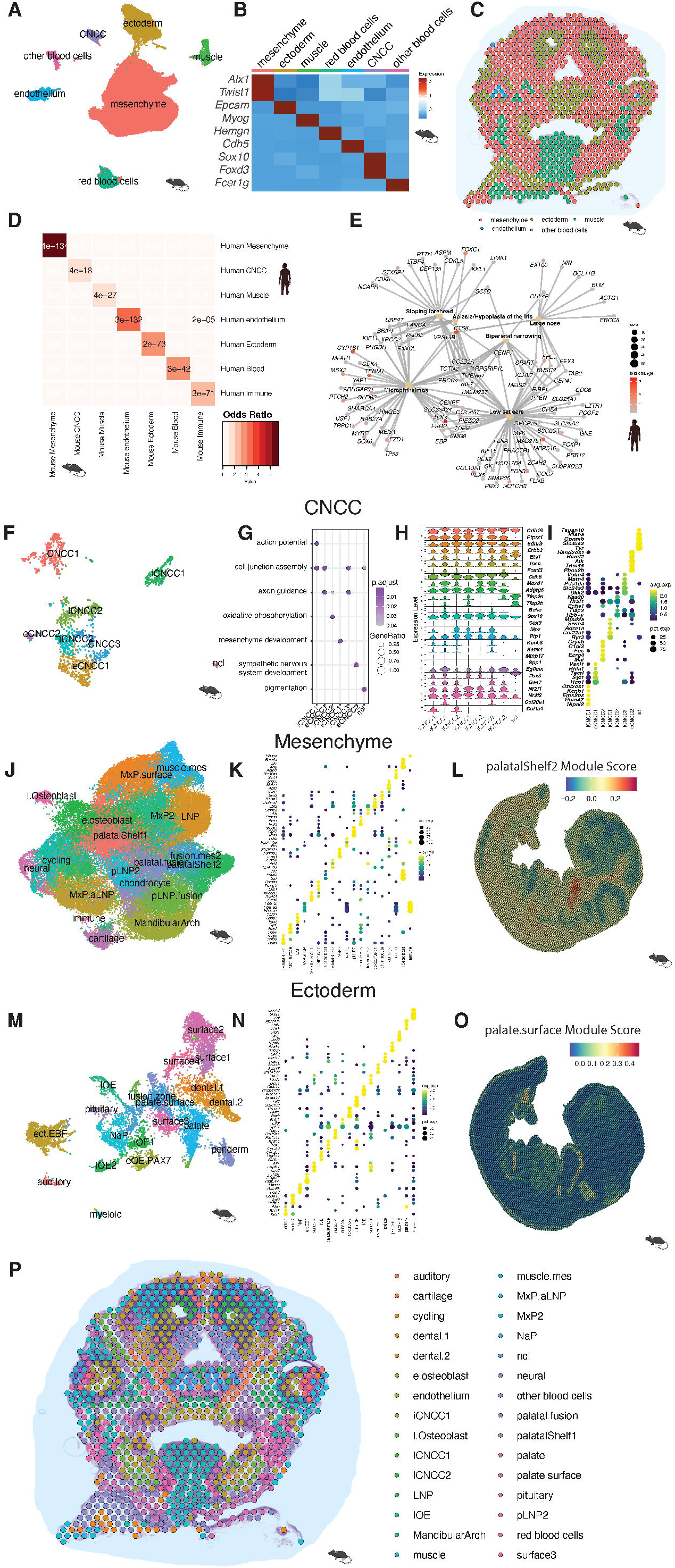
Single-nucleus gene expression in the developing mouse face. A). UMAP projection of all cells profiled by this study and combined with published studies. Major cell types are indicated. B) Heatmap of expression for indicated marker genes across each cluster. C). Spatial prediction of major cell types across E15.5 craniofacial section from Pina et al 2023^23^. D). Heatmap of sharing of marker genes between each major cell type in human and mouse. (P-values calculated by GeneOverlap R package). E). Network plot of human specific mesenchymal markers related to selected ontology categories. Shading of individual gene nodes based on fold change in expression of cells in the mesenchymal main cell type versus all other cell types. F). UMAP projection of subclustered CNCC cells from mouse. G). Gene ontology enrichments for indicated categories across each CNCC subcluster. H). Violin gene expression plots across CNCC subcluster for neural crest gene orthologous to human genes plotted in Fig. 3C. I). Average and percent expression for the top 5 marker genes for each of the mouse CNCC subclusters. J). UMAP projection of subclustered mesenchymal cells from mouse. K). Average and percent expression for the top 5 marker genes for each of the mouse mesenchymal subclusters. L). Spatial feature plot for module scores of top 100 marker genes of the PalatalShelf2 subcluster on a section of a mouse E11.5 embryo^43^. M). UMAP projection of subclustered ectodermal cells from mouse. N). Average and percent expression for the top 5 marker genes for each of the mouse ectodermal subclusters. O). Spatial feature plot for module scores of top 100 marker genes of the palate surface subcluster on a section of a mouse E11.5 embryo^43^. P). Spatial predictions of selected craniofacial subtypes on E15.5 craniofacial section from Pina et al 2023^23^.

### Species-specific differences in marker gene expression during human and mouse craniofacial development

While the overall craniofacial cell types and major gene expression patterns were shared between species, our analysis revealed hundreds of marker genes that were only called in a single species (Supplemental Tables 19-21). The largest fraction of species biased calls was observed for CNCCs. As expected, the shared CNCC markers were enriched for functions related to gliogenesis and nervous system development. However, the human-biased markers were biased toward genes related to ribosome biogenesis and cytoplasmic translation while mouse-biased markers were enriched for genes with functions related to oxidative phosphorylation and the electron transport chain (Figure S10A-D). When we examined the mesenchyme cluster, we found the shared markers were enriched for morphogenesis and differentiation programs for mesenchymal derived cell tissues as expected. However, human-biased markers were enriched generally for functions related to DNA replication and cell cycle while mouse-biased markers were enriched for only a few categories primarily related to MAPK signaling pathways (Figure S11 A-E). When we examined the human disease phenotypes enriched for each of these gene sets, we found general craniofacial abnormalities and isolated cleft palate among conserved genes. Mouse-biased mesenchymal markers were enriched exclusively for multiple seizure disorders. Human-biased mesenchymal markers were enriched for a number of craniofacial related phenotypes including microphthalmos and low set ears and exclusively for sloping forehead, large nose, and biparietal narrowing (Figure S11D). Given these phenotypes, genes driving these enrichments could be significant contributors to differences in skull shape, size, and function between human and mice. When we inspected these categories, we found genes with the highest levels of specificity for human mesenchyme included *ALX3*, *CTSK*, *CYP1B1*, *FOXC1*, *MAB21L1*, *MSX2*, and *TENM1* (Fig. 4E).

### Leveraging mouse craniofacial cell-type annotations to identify human craniofacial subtypes

Having demonstrated that major cell types, including the CNCCs, could be readily identified in both species and showed significant conservation of gene expression, we reasoned we could leverage the substantial annotation resources that have been generated for mouse to identify human cell subtypes. To achieve this, we focused on the major cell types that have been extensively subclustered and characterized in previous publications^12,23,74^, mesenchyme and ectoderm, as well as the novel populations of CNCCs we have identified here. When we performed subclustering of mouse CNCCs, we identified 8 distinct subtypes (Fig. 4F). These had a very similar arrangement in UMAP space compared to the subclusters we identified in the human CNCCs (Fig. 3A). When we examined functional enrichment of marker genes of each of these subclusters we found similar results as in human, including a clear population of melanocytes (Fig. 4G, Supplemental Table 22). Examination of the same neural crest markers as in human CNCC subtypes revealed very similar patterns of expression (Fig. 4H). We observed that *Sox10* and *Nr2f2* were expressed across all the subtypes as well as a similar trend in variable expression of *Foxd3*, *Pax3*, *Tfap2a*, and *Tfap2b* and across subtypes. When we inspected the markers for each of these subtypes, we found many of the same genes as in human subtypes including *Alk*, *Alx4*, *Crabp1*, and *Hand2* (Fig. 4I, Supplemental Table 23). When we reprocessed mouse E11.5 spatial transcriptomic data^43^ in a similar fashion to the human CS13 data, we found very similar patterns of expression for many of the human CNCC markers in mouse tissues (Figure S12A). Examination of module scores calculated from the mouse CNCC subtypes also revealed similar patterns across the mouse embryo as observed for human (Figure S12B). To attempt to identify the orthologous CNCC subtypes across species we compared sharing of orthologous marker genes much as we did with the main cell types. When we examined a confusion matrix of comparisons of cell types we found the highest similarity amongst CNCC subclusters human iCNCC4 and mouse iCNCC2 as well as human cnl4 and mouse cnl, the putative melanocyte clusters (Fig. S13A). The additional cnl clusters in human showed variable similarity to mouse and could reflect heterochrony, primate cell states not present in rodents, or the more genetically diverse human samples profiled.

Having demonstrated that even in the potentially least conserved cell type that we could readily identify shared subtypes across species we then turned to the other major cell types, mesenchyme and ectoderm. We subclustered the large mouse mesenchyme cluster and identified 19 subclusters across the mouse timeseries. Using a combination of gene ontology enrichments of marker genes and previous annotations of mouse craniofacial single cell and spatial transcriptomics we assigned functional and/or positional labels to each cluster (Fig. 4J, Supplemental Tables 22 and 24-27). For example, the well-established lateral nasal process (LNP) marker *Pax7*^12,75–77^ and the osteoblast marker Sp7^78,79^ were used to define respective clusters. In the case of osteoblasts we observed two clusters expressing similar markers but were biased in cells from different stages of development, thus we further refined these as early and late osteoblasts (Figure 4K). Two small clusters clearly represented contaminating blood derived cells or neuronal-like populations while one additional cluster could not be readily identified but had many markers associated with rapidly cycling cells (Fig. 4K). When we examined the mouse E11.5 spatial transcriptomics data we had reprocessed above, we found good concordance between marker gene expression and generalized localization in the embryo (Figure S14A). In contrast to both the human and mouse CNCC analysis, projection of modules scores for mouse mesenchymal clusters readily identified specific regions of the developing craniofacial structures that corresponded well to labels we had assigned them (Fig. 4L and S14B).

We performed identical analyses for the ectodermal cluster revealing an additional 19 subclusters (Fig. 4M). Applying the same analysis of marker genes from the literature, gene ontology enrichments, and expression in mouse single cell transcriptomics data we annotated each of these clusters with functional and spatial labels (Supplementals Tables 22 and 24-27). We identified highly specific ectodermal populations like periderm marked by *Gabrp*^12,80,81^ , cells that will form structures of the inner ear marked by Oc90 (Zhao et al 2007, Wang PNAS 1998), palate ectoderm identified by *Foxe1*^82^, and the putative pituitary marked with *Lhx3* and *Lhx4*^83,84^ among others (Fig 4N). As with the mesenchyme, we found the spatially resolved expression of marker genes corresponded well to expected positions of the mouse embryo (Figure S15A). Module score calculations for each subtype resulted in refined spatial identification of subtypes that matched the labels and expected positions well (Fig. 4O and S15B).

While the module score analysis is indicative of the cell types and spatial locations of the labels we applied, they are calculated independently of any other cell types. We therefore sought to predict what are the dominant cell types in specific locations based on spatial transcriptomics data we had not used for any of the previous analysis. When we projected top spatial predictions for 20 of the subtypes identified across previously published E15.5 mouse head data^23,85^ we found very good concordance for cell type labels and known anatomical features (Figure 4P). Overall, the analyses performed here confirmed the identities of multiple cell types across the development of mouse craniofacial tissues. Moreover, the demonstration of conserved marker genes provides a framework for transferring cell type labels to subclusters identified in human data as well as putative spatial inferences from data that originally lacked that information. We explore the subtype identifications in human data below.

### Characterization of mesenchymal cell subtypes

When we subclustered the large number of mesenchymal cells, we identified 22 subtype clusters. Using the same confusion matrix-based approach from above based on orthologous gene expression in mesenchymal subtypes, we assigned cell type and/or functional labels to each of the human clusters (Fig 5A and S13B). In some cases, multiple human clusters correlated well with a single mouse cluster and were labelled as separate populations (e.g., mouse mandibular arch and human mandibular arch 1-3). The most abundant cell types were obtained from the mandibular arch and the maxillary process and were well represented from each of the timepoints. Some of the transient structures like the lateral nasal process and cells labeled as early osteoblasts were biased towards early timepoints, while later forming cell types and structures such as cartilage and palatal shelves were dominated by cells derived from CS20 samples (Fig. 5B). When we examined marker genes identifying each of these clusters we found many transcription factors including *BARX1*, *MSX1*, and *MSX2* in the maxillary process population 2 cluster (MxP2); *SHOX* in mandibular arch 1 (arch1); *PAX7* in lateral nasal process population 2 (LNP2); *SPX* in palatal fusion zone population 1 (palatal.fusion.1); *HAND1* in mandibular arch population 3 (arch3); *HOXA3*,*B3*, and *D3* in fusion mesenchyme population 1 (fusion.mes.1); *MKX* in palatal shelf population 1.1 (palatal.shelf.1.1); and *TBXT* in cartilage population 2 (cartilage.2) among many others (Fig. 5C).

**Figure 5.**
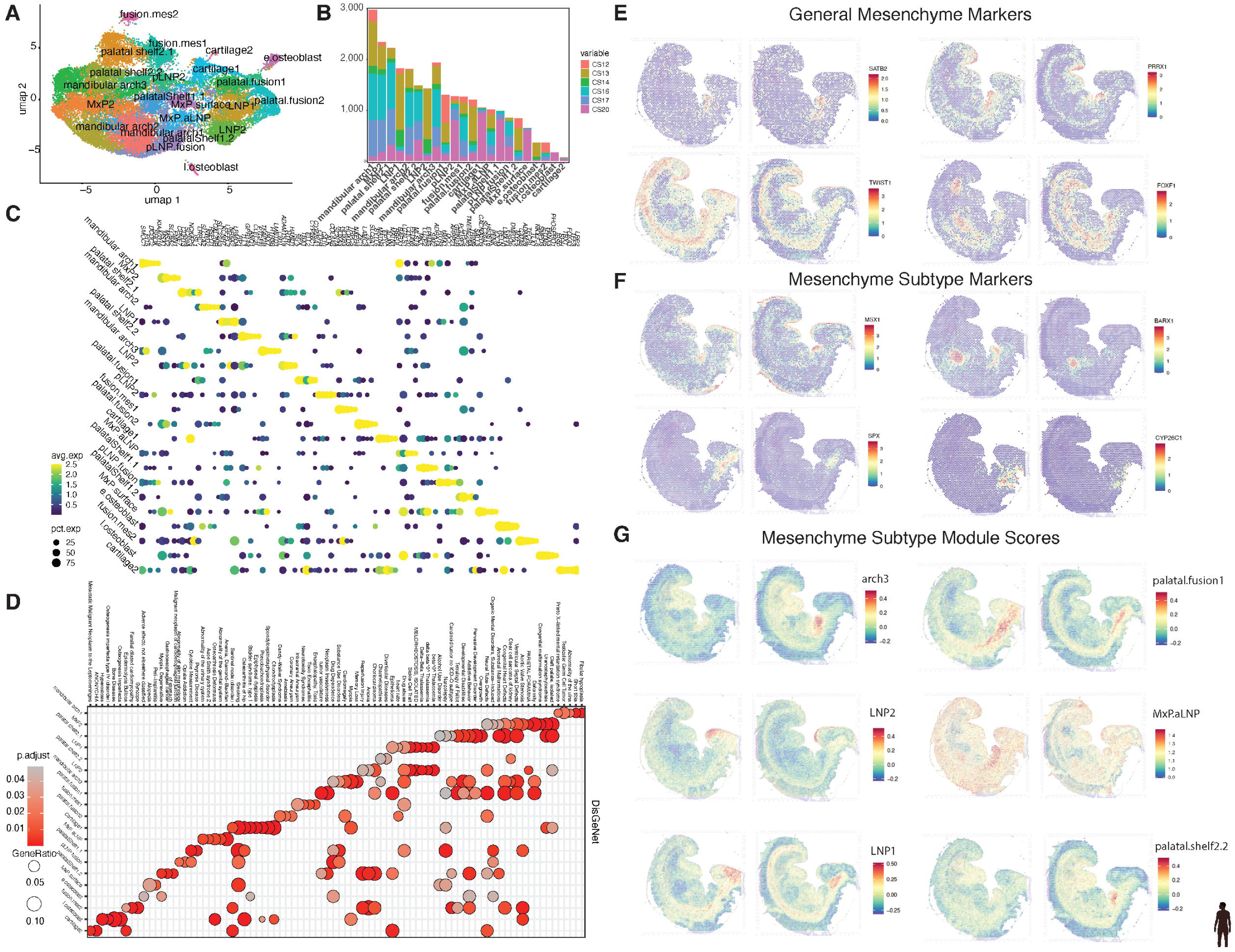
Identification of mesenchymal subtypes in human craniofacial development. A). UMAP projection of subclustered mesenchymal main cell type. Subtype labels based on transfer of mouse mesenchymal subtypes to human. B). Contribution of cells from each CS timepoint to each mesenchymal subcluster. C) Average and percent expression for the top 5 marker genes for each of the mesenchymal subclusters. D). Disease ontology enrichments for each of the indicated mesenchymal subcluster. E.) Gene expression spatial feature plot for indicated mesenchymal marker genes in two sections from a CS13 human embryo. F). Gene expression spatial feature plot for indicated mesenchymal subtype marker genes in same sections as E. G). Spatial feature plot for modules scores calculated from top 100 marker genes from indicated mesenchymal subtype.

Gene ontology analysis revealed many biological processes, cellular component, and molecular process categories that were relevant for these subtypes (Figure S16). For example, cartilage1 and cartilage2 were differentially enriched for hyaluronic acid and frizzled binding respectively. Cartilage 1 is primarily found in CS20 samples suggesting these are distinct stages of cartilage development. Early osteoblast markers were enriched in pathways regulating pluripotency while late osteoblast markers were enriched for PI3K-AKT signaling and parathyroid hormone response. The more regional based annotations shared many of the same functional enrichments suggesting the same underlying processes were active in these cell types. However, the maxillary process / anterior lateral nasal process derived cells (MxP.aLNP) likely from near the lambdoid junction^86^showed substantially higher expression of many genes related to ribosome production and cytoplasmic translation. Examination of disease enrichments across cluster marker genes revealed some tissue-specific disorders like Osteogenesis imperfecta in late osteoblasts and epiphyseal dysplasia in cartilage 1. Enrichment for genes related to isolated cleft palate were found in several clusters including MxP2, palatal.shelf2.1, palatal.shelf.2.2, and cartilage 1 (Fig 5D).

While the gene ontology analysis confirmed the labeling of some specific cell types, the more positional types remained less clear. To address this, we again turned to the CS13 human spatial transcriptomics data. When we examined some of the markers that defined the mesenchyme versus other cell types, such as *TWIST1* and *PRRX1*, we found fairly broad expression across the embryo with some bias toward the craniofacial region. Other markers like *SATB2* were much more regionally restricted and potentially specifically mark craniofacial mesenchyme versus other types (Fig. 5E). When we examined some of the subtype marker genes, we found much more regionalized expression. *MSX1* was found near many surface locations with a bias toward the head. *BARX1* was rather specifically localized in the general region of the pharyngeal arches and the developing stomach. *SPX* and *CYP26C1* were both restricted to the head region of the embryo at this stage (Fig. 5F). As was observed in mouse, we found much more regionalized signals from module scores for each subtype. The mandibular arch clusters were clearly enriched in the pharyngeal arch region of the CS13 embryo and biased toward the more anterior portion of this region. The lateral nasal process clusters were enriched in distinct areas of the head with LNP1 being more posterior and LNP1 being more anterior. Other subtypes like MxP2 and palatal.shelf2.2 showed good spatial concordance with the labels that had been assigned (Fig. 5G).

### Characterization of ectodermal cell subtypes

We then turned to the ectodermal cluster to identify potential subtypes. Using the same basic approach as the mesenchyme, we identified 22 distinct ectodermal clusters (Fig. 6A). Transferring of mouse labels (Figure S13C) revealed cells that would give rise to specific ectodermal-derived organs like the pituitary and thyroid, structures of the inner ear (auditory1-3), and surfaces of several structures including periderm (Fig. 6A). As with mesenchymal subclusters, many of the ectodermal subclusters annotated as early versus late had biased sample contributions (Fig. 6B). Amongst marker genes of ectodermal subtypes, transcription factors were again prominent. *LHX3*, *SIX6*, and *PITX2* were most strongly expressed in the pituitary; *GATA6* marked the palate subtype; the nasal placode (NaP) was identified by *SP8* and *FEZF1*; auditory subtypes 1-3 were marked by *SALL3*, *GRIN2A*, and *SP9* respectively; *EBF1*, *EBF2*, and *EBF3* in a single ectodermal subtype (ect.EBF); and *TBX18* marked a putative fusion zone cluster among others (Fig 6C).

**Figure 6.**
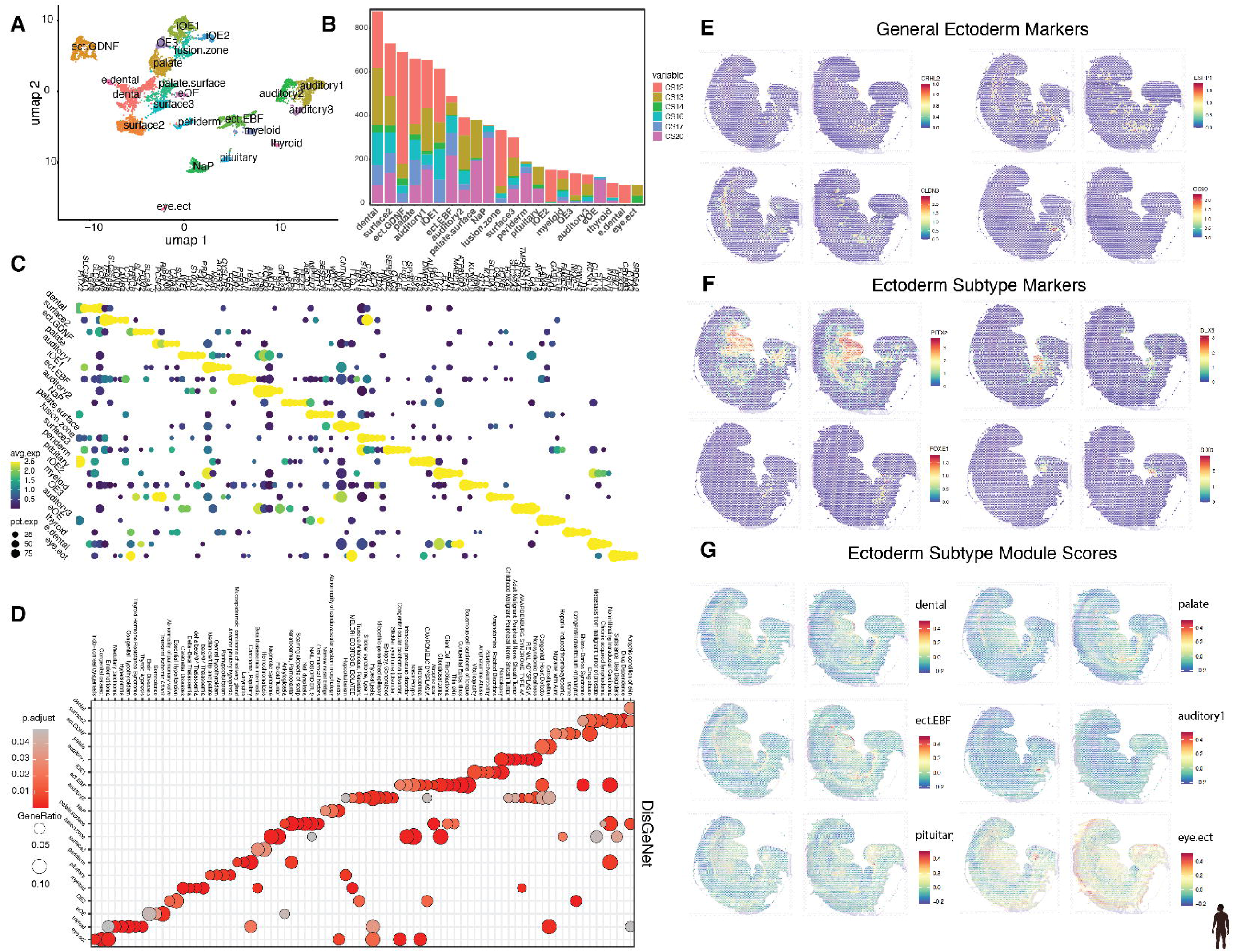
Identification of ectodermal subtypes in human craniofacial development. A). UMAP projection of subclustered ectodermal main cell type. Subtype labels based on transfer of mouse ectodermal subtypes to human. B). Contribution of cells from each CS timepoint to each ectodermal subcluster. C) Average and percent expression for the top 5 marker genes for each of the ectodermal subclusters. D). Disease ontology enrichments for each of the indicated ectodermal subcluster. E.) Gene expression spatial feature plot for indicated ectodermal marker genes in two sections from a CS13 human embryo. F). Gene expression spatial feature plot for indicated ectodermal subtype marker genes in same sections as E. G). Spatial feature plot for modules scores calculated from top 100 marker genes from indicated ectodermal subtype.

Consistent with our findings for CNCCs and mesenchyme, gene ontology analysis revealed many biological processes, cellular component, and molecular process categories that were relevant for ectodermal subtypes (Figure S17). For example, markers for all three auditory subtypes were enriched for terms related to inner ear morphogenesis and development; genes biased for eye ectoderm were enriched for structural components of the lens; periderm marker genes were associated with the cornified envelope and skin development; markers for the thyroid cluster were enriched for thyroid hormone synthesis; and the markers of the pituitary were associated with pituitary gland development. The less specific clusters such as ectodermal surface clusters were enriched for a variety of categories suggesting they might be more regionally distinct cell states. In particular, surface3 marker genes were biased for oxidative phosphorylation and cytoplasmic translation compared to other ectodermal subtypes (Figure S17A-E). Examination of enriched human diseases revealed many tissue- or region-specific disorders including aniridia in the NaP cluster; nonsyndromic deafness in auditory clusters 1 and 2; thyroid agenesis for the thyroid cluster; anterior pituitary hypoplasia for the pituitary cluster; and congenital cataracts in the eye ectodermal cluster. Interestingly, median cleft lip and palate was only enriched in the pituitary cluster. Lastly marker genes of the ectodermal cluster expressing high levels of *EBF* genes (ect.EBF) were enriched for the largest number of disease categories suggesting this might be a particularly disease relevant cell type or state (Figure 6D).

Examination of overall ectodermal markers revealed relatively restricted expression to various surfaces in the human spatial transcriptomics data. One notable exception being OC90 that was strongly expressed in the location of the putative inner ear (Fig. 6E). Subtype markers also showed generally restricted expression particularly for *DLX5, FOXE1,* and *SIX6*. *PITX2* was expressed in multiple putative fusion locations in the head but also strongly in the hindlimb (Fig. 6F). Markers of the ect.EBF subcluster, *EBF2* and *EBF3*, were biased in expression toward the head and pharyngeal arches of the CS13 human embryo. When we examined the spatial expression for both human CS13 and mouse E11.5, we found qualitatively different patterns of expression in the craniofacial regions corroborating our previous finding (Fig. S18). When we inspected module scores of each subtype, we observed exquisitely specific localization for some clusters like pituitary and auditory. Other clusters were generally enriched at surfaces of the pharyngeal arches and the putative esophagus (Fig. 6G).

### Cell-type specific enrichment of genes and variants linked to orofacial abnormalities and normal facial variation

The analysis above demonstrated strong concordance between human and mouse cell types and subtypes, showed coherent functional and disease enrichments across these cell types, and revealed spatial enrichments consistent with functions and expected anatomical locations. The strong support of our labelling of cell types across human craniofacial development, gave us the opportunity to interrogate the cell type-specific expression profiles for enrichment of craniofacial related genetic signals. The genetic contributions of common variants to many aspects of craniofacial variation have been studied in multiple populations based on frontal and profile photographs^87,88^. However, the cell types and embryonic landmarks that drive these differences are currently unknown. To address this issue, we first processed the genome-wide summary statistics^87,88^ for each craniofacial landmark measurement with at least one genome-wide significant association using the linkage disequilibrium aware approach Multi-marker Analysis of GenoMic Annotation (MAGMA^89^. We then calculated expression weighted cell type enrichments^90^ (EWCE) across all the cell types identified in our study using MAGMA-Celltyping^91^. We observed distinct patterns of cell type enrichment related to different sections of the face. We found that profile landmark measures related to soft tissues including multiple measures of lip thickness and shape, ear size, and nose shape were enriched primarily in ectodermal subtypes. Frontal landmark measures related to aspects of these same portions of the face such as the distance of the outer edge of the eye to the nasion (ExR-N) showed similar patterns of ectodermal enrichment. Measures likely to be influenced by bone or cartilage structure such as jaw, chin, and brow protrusion as well the positioning of the eyes relative to the base of the nose (EnL-Sn) and the mouth (ExR-ChR) were enriched primarily amongst mesenchymal subtypes (Fig. 7A). Amongst mesenchymal cell types, the mandibular arch, palatal shelf, and maxillary process fusion zone subclusters had largest number of significant enrichments for facial shape. The fusion zone cluster and surprisingly the pituitary cluster had the largest number of significant enrichments amongst ectodermal subtypes. Interestingly, while CNCCs certainly give rise to many of the downstream cell types and tissues, we found relatively few shape associations for CNCC subtypes. Overall, this analysis suggests specific cell subtypes contribute differentially to individual facial differences and suggest these effects begin to manifest very early in human development.

**Figure 7.**
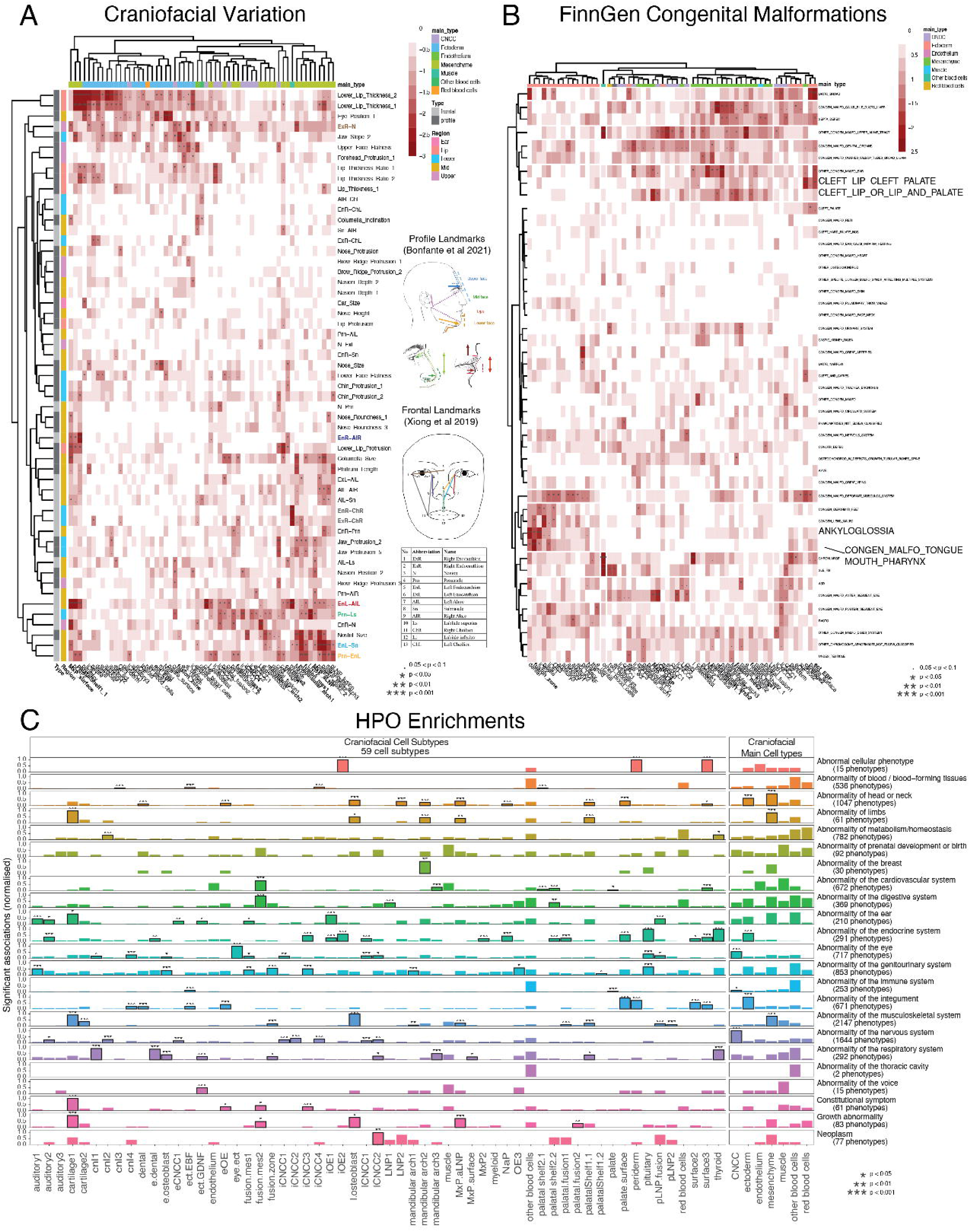
Enrichment of common variation associate with facial shape differences and congenital abnormality risk. A). Clustered heatmap of significance values calculated by MAGMA Celltyping^91^ for each facial variation trait and cell subtype. Profile landmark diagram adapted from Bonfante et al 2021^88^. Frontal landmark diagram adapted from Xiong et al 2019^87^. Colors along top of heatmap indicate main cell type classification. Shaded gray indicators along left of heatmap indicate study origin. Colors along left of heatmap indicate general region of the face each landmark is located. Hyphenated trait measures are obtained from Xiong et al 2019 and combinatorial code is indicated in coded legend (e.g., EnL-AIL indicates landmark segment 5 to 7). Descriptive named traits obtained from Bonfante et al 2021^88^. Levels of significance indicated by asterisks or period according to figure. B). Clustered heatmap of significance values calculated by MAGMA Celltyping^91^ for each congenital abnormality or disease and cell subtype. Colors along top of heatmap indicate main cell type classification. Levels of significance indicated by asterisks or period according to figure. C) Barplot showing the number of enriched human phenotypes (max-normalized from 0-1 within each branch) for main cell types and subtypes as calculated by MSTExplorer::run_phenomix. Significance of the proportion tests, testing for disproportionate numbers of phenotype enrichments for a given cell type within a given HPO branch, is denoted with asterisks (FDR<0.001=***, FDR<0.01=**, FDR<0.05=*) as well as black outlines around the bars.

We next turned to studies of the genetic underpinnings of craniofacial abnormalities. In particular, there have been dozens of genetic associations identified for risk for orofacial clefting in multiple populations^92–106^. However, the cell types that potentially influence risk for clefting have not been identified in human development. While orofacial clefting has been examined extensively using a variety of approaches, these studies have been performed in many different populations, making cross-study comparisons challenging ^93,107–115^. Moreover, to identify true positive signals for cell type enrichments diseases that are not expected to be related to craniofacial cell types examined in the same population are needed as negative controls. To mitigate these issues, we turned to genome wide association studies that have been systematically performed on a large cohort of Finnish ancestry^116^. From this resource we selected all studies annotated as a congenital abnormality by FinnGen with at least one genome-wide significant association (n = 45) as well as two immune related diseases, Crohn’s disease and systemic lupus erythematous (SLE), that we have used as negative controls in our previous studies^28,117^. When we analyzed these GWAS using the same approach as for facial variation we found similar partitioning of enrichment between specific classes of cell types (Fig 7B). We found that ectodermal cell subtypes were enriched for cleft lip with cleft palate (palate.surface), ankyloglossia, and other congenital malformation of the tongue and mouth (dental, fusion.zone). Mesenchymal subtypes were enriched for cleft lip or lip and palate (MxP_aLNP, mandibular arch 3, fusion mesenchyme subcluster 1 and 2), other congenital malformations of the ear (multiple palatal shelf subtypes), and congenital malformation of the musculoskeletal system (cartilage2) among others. CNCC subtypes were most consistently enriched for other congenital malformations of the upper alimentary tract. The immune cluster was most significantly enriched for Crohn’s and SLE. Many other congenital abnormalities showed no significant enrichments for any craniofacial cell types demonstrating the specificity of our analyses. A few of these cell types, MxP_aLNP and ect_EBF in particular, were associated with both craniofacial disease and normal facial variation. These findings suggest that some cell types are contributors to both facial shape as well as risk for clefting. These results also point to underlying differences in how clefting phenotypes are categorized which are then in turn related to different subtypes of mesenchyme and ectoderm.

Our results from the marker gene ontology enrichments and common variant associations point to relevant craniofacial disease and phenotype enrichments for specific craniofacial cell types. However, it is unclear if these cell types might be generally informative for other human phenotypes. We posited that integrating continuous expression patterns instead of just binary marker gene identity may reveal additional associations. To address this, we employed a systematic examination of the entire Human Phenotype Ontology (HPO)^118,119^ (Fig 7C). As expected, the immune cluster was systematically enriched for 90 of 253 phenotypes related to abnormality of the immune system. The red blood cell cluster was enriched for terms related to abnormality of metabolism and homeostasis (60 of 782 phenotypes). Both these cell types were enriched for phenotypes related to abnormalities of blood and blood-forming tissues (150 and 75 of 536 respectively). The endothelium cluster was enriched for abnormality of the cardiovascular system (60 of 672 phenotypes).

Amongst ectodermal subtypes we found the eye subcluster was strongly enriched for phenotypes related to abnormalities of the eye (75 of 717). Periderm, palate surface, and surface 2 and 3 subtypes were enriched for abnormalities of the integument. As expected, the pituitary and thyroid subtypes were associated with abnormalities of the endocrine system. Surprisingly many of the ectodermal subtypes were enriched for phenotypes related to abnormalities of the respiratory and genitourinary systems. Among the mesenchymal subclusters, many were enriched for abnormalities of the head and neck. The cartilage1 cluster showed the most diverse enrichments including phenotypes related to growth abnormalities and abnormalities of the musculoskeletal system, ear, and limb. The main CNCC cluster was enriched for abnormalities of the nervous system, driven by most of the CNCC subclusters with the exception of the cnl1,3, and 4 subclusters. Surprisingly the specialized ectodermal subtype ect.GDNF was significantly associated with abnormalities of the voice. Together these results suggest that some subtypes we identified are not specific to the head and are more general states like cartilage. Moreover, this analysis revealed that while no major cell types were enriched for neoplasms, late CNCCs employ gene expression programs that likely trigger overgrowth.

### Differential enrichment of curated gene lists revealed distinct disease risk and role in skull shape and/or function across hominid evolution

Thus far our analysis of the craniofacial cell types has leveraged annotated ontology categories and common variant associations. Other gene lists that are not part of these systematic ontology databases and potentially of use to the craniofacial field have not been interrogated. To address this were assembled multiple gene lists relevant for orofacial clefting including those compiled by CleftGeneDB^120^, genes co-expressed in important gene modules or prioritized for craniofacial disease in our recent work^13^, and genes with distinct classes of *de novo* mutations (synonymous vs protein altering) in orofacial cleft trios sequenced as part of the Gabriella Miller Kids First program ^121,122^ and CPSeq Studies^123^. We also curated genes at the extremes of tolerance to loss of function mutations in otherwise healthy populations that have been suggested to be enriched or depleted of disease relevant genes^124^. Lastly given our findings for common facial variation across humans, we wondered whether genes potentially regulated by Neanderthal derived sequences might have craniofacial cell type specific enrichments. As a control for this evolutionary analysis, we included genes near human accelerated regions, which have been reported to be enriched in neuronal related functions and expression^125–128^.

With these lists in hand, we again employed the expression weighted cell type enrichment approach. We found that the CleftGeneDB, craniofacial black co-expression modules, and our prioritized gene lists showed similar patterns of significant enrichments in mesenchymal subtypes including multiple clusters related to the maxillary process, palatal shelves, and lateral nasal process (Fig. 8A). Relatively few ectodermal and CNCC subtypes were enriched for these gene lists. The genes identified by gnomAD to have the least tolerance for loss of function mutations (LOUEF decile 1) were significantly enriched in many different subtypes identified by our analysis. In particular, MxP.aLNP cluster showed the most significant enrichment. This contrasted with those genes with the most tolerance for loss of function mutations (LOUEF decile 9) that showed few enrichments and were generally non-overlapping with the LOUEF decile 1 enrichments (Fig. 8A).

**Figure 8.**
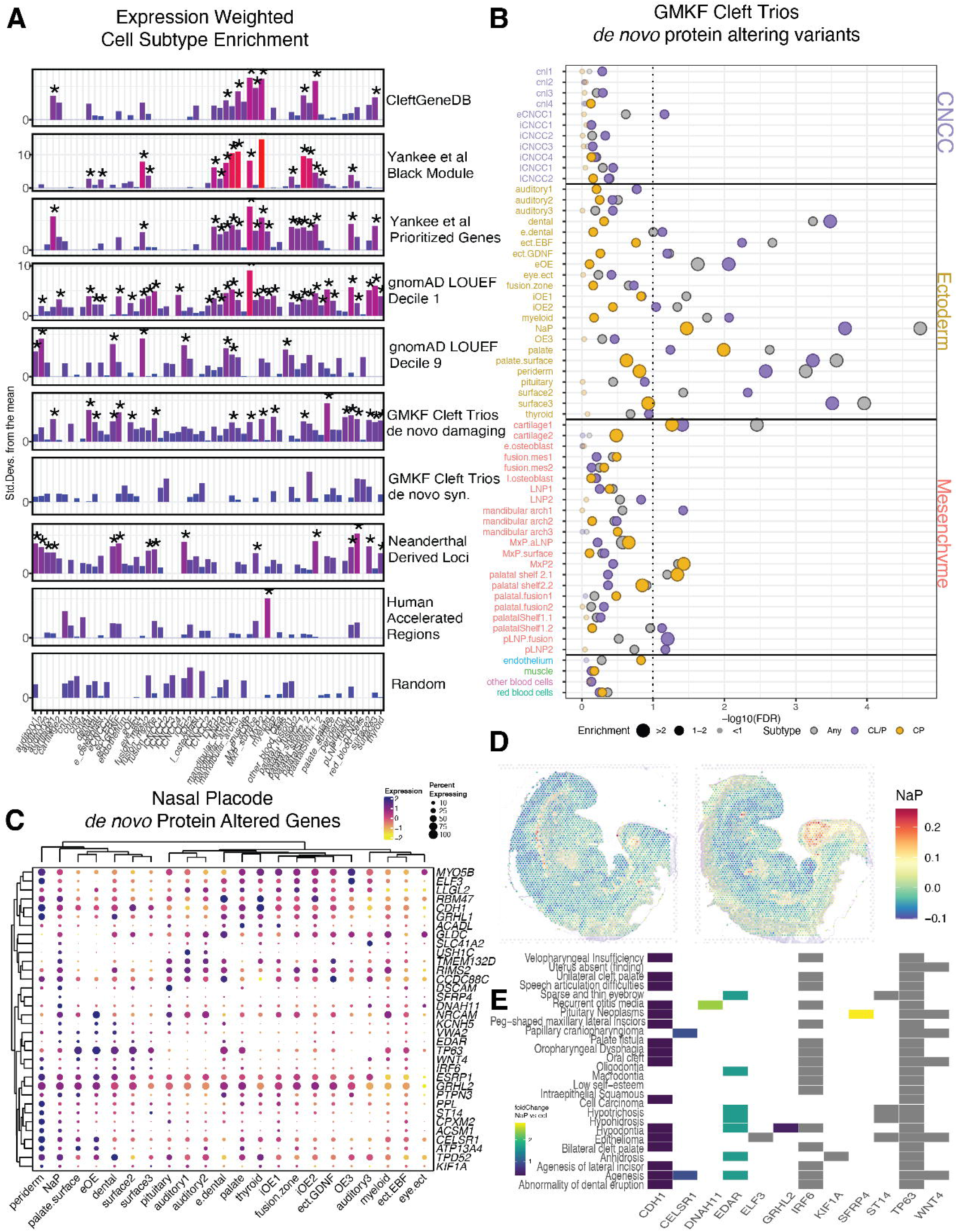
Genes associated with orofacial clefting, constraint in human populations, and Neanderthal introgression show distinct cell subtype enrichments. A). Bar plot of standard deviations from the mean of bootstrapping tests performed by EWCE method^90^ for each indicated gene list and cell subtype. Asterisks indicate significant subtype enrichments corrected for number of gene lists and cell subtypes performed for entire figure. B). Bubbleplot of -log10 transformed significance and fold enrichment values for each cell subtype from denovolzeR^129^ analysis of protein *damaging de novo* variation in orofacial cleft trios from the Gabriella Miller Kids First program^121,122^. Colored circles indicate variants identified in whole cohort (Any), in cleft lip with cleft palate probands (CL/P), or cleft palate only probands (CP). Cell subtypes are clustered by main cell type. C) Average and percent expression across all ectodermal subtypes of genes identified in nasal placode subtype *with de novo* protein damaging mutations for B. D). Spatial feature plot of modules scores calculated from top 100 marker genes from nasal placode ectodermal subtype on CS13 human embryo. E). Heatmap of fold differences in expression of each indicated gene in nasal placode subtype versus all other ectodermal cells. Presence or absence of box indicates membership in indicated disease ontology category indicated as significantly enriched in nasal placode marker genes.

When we analyzed the genes near Neanderthal derived sequences, we found patterns of cell type enrichment distinct from the more disease-focused lists described above. The strongest enrichment was observed in pLNP2 mesenchyme subtype. We identified the specialized ect.EBF and ect.GDNF clusters, two fusion mesenchyme subtypes, and cartilage1. Interestingly all three auditory types were significantly enriched in this analysis. This was contrasted by only a single cell type identified when examining HAR associated genes, consistent with their previously published association with brain cell types and neuronal function ^125–127^. We found no consistent, significant enrichments from any of our randomly selected gene lists across the cell types in questions. We also found no enrichments for the red blood cells across any of these gene lists, and only the gnomAD unconstrained genes for the immune cell types (Fig. 8A).

We then turned to recently identified *de novo* variants from orofacial clefting trios from the Gabriella Miller Kids First program^121,122^ and CPSeq studies^123^. We found no enrichment in any cell types for genes affected by *de novo* synonymous variants. We found no enrichment in any cell types for genes affected by *de novo* synonymous variants. However, we identified multiple cell types that strongly express genes with *de novo* protein altering variants (Fig. 8A). Palate ectoderm showed the strongest enrichment from this analysis, a cell type that was not enriched for any of the community curated gene lists related to clefting nor our previous prioritized genes^13^. Multiple other ectodermal cell types were also identified as enriched including multiple surface ectodermal subtypes, specialized ectoderm ect.EBF and ect.GDNF, fusion zone ectoderm, and the nasal placode (NaP). Fewer mesenchymal cell type enrichments were observed but identified the MxP.aLNP and others related the lateral nasal process (pLNP2, pLNP.fusion).

These findings suggest that current disease associations have been biased for genes expressed in the mesenchyme and that many genes expressed in ectodermal subtypes are also substantial contributors to clefting risk. To explore this concept further we wondered whether not only the number of genes, but the total number of *de novo* variants observed in genes might reveal additional disease associations. When we applied a computational framework that examines gene lists for excess *de novo* mutational load^129,130^ we largely confirmed the findings from the EWCE analysis. We identified 22 clusters for which a least one phenotype was significantly enriched using a Benjamin-Hochberg false discovery rate of <10% (Fig. 8B and Supplemental Table 30). These included 17 for all trios with OFCs, 19 for trios with CLP, and 5 for trios with CP. We identified 14 significant enrichments across ectodermal cell types (n=22), 7 enrichments from mesenchymal cell types (n=22) and a single CNCC subtype (n=11). Only 3 clusters were significantly enriched in all three categories (NaP, palate, and cartilage1), whereas there were 12 shared between all OFCs and CL/P and 2 shared between all OFCs and CP. We also found 5 clusters that were only significant in the CL/P group and 1 that was only significant in the full cohort. No significant findings were observed for endothelium, muscle, red blood cell, or immune cell types in our data.

For the ectodermal subtypes we identified strongest enrichment *for de novo* variants identified in the whole cohort and those probands with cleft lip with cleft palate (CL/P) in the nasal placode, surface3, and palate.surface. We only identified significant enrichment *of de novo* variants from cleft palate only probands (CP) in the nasal placode and palate ectoderm. While fewer significant enrichments were observed for mesenchymal subtypes, we found cartilage1 was enriched for all analyses performed. Interestingly several subtypes were biased toward significant enrichment related to CP vs CL/P. For instance, MxP2 and palatal shelf 2.1 were enriched for the former while pLNP2 and pLNP.fusion for the latter (Fig.8B).

To explore the genes driving these enrichments we examined the genes with *de novo* damaging variants that were markers for the nasal placode, the most significantly enriched subtype across our analysis. As expected, these genes were all expressed in the NaP cells, but were frequently expressed in many other types of ectoderm to varying degrees (Fig. 8C). In particular, a high degree of sharing of expression was observed with periderm and multiple surface subtypes, including genes previously implicated in orofacial clefting like *TP63*, *IRF6*, and *CDH1*. Among the *de novo* damaged genes those with the most biased expression in NaP were *SFRP4* and *DNAH11*. When we examined the localization of the NaP cells on the spatial transcriptomics data, we found discrete localization at the putative frontonasal and maxillary processes (Fig 8D). Finally, when we examined these genes for known disease enrichments, we found enrichment for various types of clefting and craniofacial abnormalities (Fig 8E). These were driven largely by the genes listed above related to clefting. Interestingly many of the genes we identified here are expressed in similar patterns to those known disease genes, but have not been associated with many human disease phenotypes. Amongst these *SFRP4* has the highest specificity of expression across the main cell types and ectodermal cell types (Fig 8E).

Compared to the shared expression and overlapping genes between the NaP and palate clusters, the genes driving enrichment in cartilage1 were more distinct. Interestingly, although both CP and CL/P were enriched to a similar degree, the makeup of genes contributing to this signal was different. For CP, the main driver of the signal was due to *COL2A1* variants, which made up half of the observed variants, where the remainder were single gene contributions (total n=10). This gene has fairly restricted expression in the head region and presumptive somites of the CS13 embryo (Figure S19). In contrast, CL/P probands collectively were enriched within cartilage1, but there were no genes that were individually overrepresented—only *KCNH5* had multiple variants (2 of total n=18), and the rest were a single variant per gene. This enrichment highlights the importance of these cells in OFC etiology, but the difference in signal drivers may provide insight into the heterogeneity of the genetic architecture between CP and CL/P.

## Discussion

Craniofacial abnormalities are some of the most common human birth defects. Only recently have gene expression patterns active during human craniofacial development been examined^13^ . We previously showed that genes specifically or co-expressed across craniofacial development relative to other tissues were enriched for known disease-causing genes^13^. However, these analyses relied on bulk gene expression data from the developing craniofacial tissues. The face is a complex structure that is derived from multiple cell lineages like ectoderm, mesenchyme, and the specialized neural crest. These major cell types undergo differentiation to become a variety of distinct cell types that make up the face including bone, cartilage, muscle, mucosa, and vasculature. Our bulk analyses showed strong bias for gene programs expressed in human and mouse mesenchyme preventing analysis of genes in ectodermal and other less abundant cell types. While other single cell atlases from human embryonic development have been described, there were few biological replicates and relatively few cells clearly derived from craniofacial regions^24–27^. Moreover, few craniofacial centric analyses have been previously performed on such data.

Our work here attempted to address these shortcomings and concentrate on cell types that are present across many of the major milestones of human craniofacial development. In this work we profiled multiple biological replicates from six distinct stages of human craniofacial development. Across these data we identified seven major cell types present in the developing human face. Most of these, including mesenchyme, ectoderm, endothelium, blood, and immune cells, have been previously identified in mouse craniofacial development^12–16^. However, we identified two distinct clusters not described in those previous efforts or labelled as cell types not expected to exist in high levels in craniofacial tissues like glia or Schwann cells. Our thorough characterization of these clusters using curation of genes from the literature as well as extensive gene and disease ontology analyses point to these clusters being muscle progenitors and cranial neural crest. While several protocols for deriving neural crest like cells from human embryonic stem cells have been described, the primary CNCCs have remained elusive. Also, only a handful of well-known neural crest genes have been examined using immunohistochemistry in a small number of early human embryos^71^. Thus, it is unclear the complete repertoire of genes that are active in this cell type and how closely in vitro models reflect the primary gene expression patterns. Our analysis here not only established a large number of known marker genes as bona fide CNCC genes, including *FOXD3* and *SOX10*, but also identifies new genes that could be important for CNCC specification or function such as *INSC*, *ABCA8*, and *CTXND1*. Our identification of subclusters of the CNCC including putative melanocytes and the expression programs within them are likely to be useful to many researchers interested in these cell types. Moreover, identification of this exotic cell type and subtypes is not a fluke. Generation of data from mouse from similar tissues and stages and uniform process reveal these same populations. Upon close inspection of the gene ontology enrichments, other groups may have mistakenly labelled these cells as glia or Schwann cells simply because of biases in the ontology databases. Far more research has been performed on the human brain and related cell types than other parts of the body, likely resulting in many more brain related gene ontology annotations. While automated and machine learning based approaches are gaining traction for labelling of single cell atlases^131–137^, transient developmentally related cell types that are not in current databases and biases in ontology will still require close inspection and interpretation.

By generating comparable datasets from both mouse and human we had the unique opportunity to identify both shared and species-biased gene programs active in individual cell types. As expected, we found the main cell types identified in each species share the most significant amount of marker genes with the orthologous cell type in the other species. Among these, mesenchyme was the most functionally shared between human and mouse based on marker gene expression. Surprisingly, CNCC markers were the least shared between these species, even less than cells from the immune system that has been documented to have substantial differences across humans and mice^17^. This could reflect substantial functional differences in CNCC between human and mice and indicate that this cell type may be particularly labile across evolution allowing innovation of craniofacial shape as others have proposed^1,18–20,70,138^.

Although there was the largest degree of shared marker gene expression within mesenchyme, we found hundreds of differences in marker gene identity between human and mouse. While we restricted our analysis to genes with clear one-to-one orthology between these two species, some of these differences could be due to mis-annotation of orthology, substantial developmental heterochrony, or the inherit noisiness of current single nucleus gene expression data. However, by focusing on coherent gene ontologies and strongly expressed genes we identified many genes that are likely to reflect true species differences. For instance, one of the top human mesenchymal markers based on absolute and specificity of expression that was not revealed in mice was *ALX3*. Recessive mutations in human *ALX3* have been linked to frontorhiny or frontonasal dysplasia 1 (OMIM 136760)^139^, while the *Alx3*^-/-^ mouse has been reported to have no phenotype^140^. Our analysis also identified *MSX2,* to which humans have been suggested to be much more sensitive than mice to dosage of this transcription factor during craniofacial development^141^. Further analysis of all these subtypes and comparison with additional species could reveal novel functional differences as well as the core regulatory programs that are present in all vertebrates.

While we highlighted some of the species differences in major cell types that could be relevant for what human genes and diseases can be modelled in mice, our comparison framework allowed us to accurately identify subtypes of each major cell types between species. This allowed us to leverage the substantial single-cell and spatial transcriptomics resources as well cell type annotations that have been generated by many different groups^23,86,142^. By transferring functional and spatial labels for mouse cell subtypes to our human data we could add such information to data that were originally lacking. We confirmed these labels using a variety of gene and disease ontology analyses, but most convincingly by leveraging previously published spatial transcriptomics data for a CS13 human embryo^25^By using marker genes to calculate module scores across this spatial data we confirmed relevant anatomical regions from which each subtype was potentially derived. We were able to identify some exquisitely specific spatial locations for ectodermal subtypes related to the ear, eye, and pituitary. We also identified expected regionalized expression for mesenchymal subtypes putatively derived from the mandibular arch as well as important fusion zones like the lateral nasal process. Further characterization of the markers we identified in higher resolution spatial transcriptomics across multiple sections and reconstruction into a complete three-dimensional representation as has been recently described for mice will be necessary to validate these findings^143^. One of the major goals of generating such resources is to enable better understanding of human phenotypes and disease. Not only can facial abnormalities affect our capacity for communication and feeding, but the face is also one of the most defining features of each human and is intimately tied to our sense of individuality. Thus, understanding how facial shape is encoded in our genomes is of substantial general interest. In recent years coupling of two- and three-dimensional imaging approaches with large scale genotyping has enabled the discovery of common genetic variants associated with quantitative differences in many different facial landmarks^87,88,144–146^. While these variants have been shown to be enriched in regulatory regions active in the developing face, the cell types that underly facial differences were unknown. Using our highly confident cell subtype annotations, we found distinct differences in enrichments for measurements across the human face. In general, the enrichments we observed were mutually exclusive, features likely driven by mesenchyme subtypes not associated with an ectodermal subtype and vice versa. As expected, mesenchyme subtypes were associated with features that are likely driven by hard structures like bone and cartilage while ectoderm subtypes were associated with some measures that are related to soft tissue shape or thickness. The most consistent associations observed were related to variation in measures of the midface. These were significantly enriched for many mesenchyme subtypes that we annotated as derived from regions that are consistent with these effects: the maxillary process, palatal shelves, and fusion zone mesenchyme. We did not observe any subtype that contributed to all aspects of the face, nor did we observe significant subtype enrichments for all measurements. These landmarks may be driven by cell types that appear later in development or be influenced by subtle gene expression differences in many cell subtypes. While the two studies we utilized were performed in populations with distinct ancestries and yielded consistent results, it is possible that subtypes could influence facial variation differently in other populations. Further identification of genetic associations with more facial measures in a more diverse set of individuals and identification of cell types later in craniofacial development will be needed to address this issue.

Craniofacial abnormalities are among the most common birth defects in humans. The most common form of these, nonsyndromic cleft lip and/or cleft palate, is thought to occur relatively early in human development between 4 and 6 weeks^8,147,148^. Consistent with this idea, we found that variants associated with risk for orofacial clefting are enriched in regulatory sequences active in craniofacial tissues from this developmental window^28^. However, the cell types in which these variants manifest their effects were unknown. Here we used uniformly generated and processed genome-wide association data for many congenital abnormalities in the Finnish population. This population has been shown to have a high incidence of clefting with interesting geographical distributions^149^, and we reasoned would serve as an excellent test case for subtype enrichments across relevant and unrelated diseases. Indeed, we found some subtypes of both mesenchyme and ectoderm were significantly enriched for orofacial clefting or other abnormalities of mouth. We found some overlap between phenotypes and subtypes particularly related to cardiac outflow tract abnormalities consistent with the neural crest derived nature of those structures^150–154^. We found expected cell type specific enrichments for immune cells in the autoimmune related diseases that we included from this cohort, SLE and Crohn’s. We also did not observe enrichment for most subtypes in most abnormalities outside the craniofacial and cardiac structures.

Interestingly several of the enrichments we observed for subtypes were shared across the craniofacial variation and craniofacial abnormality analyses. The MxP.aLNP and ect.EBF subtypes were examples that had several significant associations in both phenotypes. This is particularly interesting as it has been speculated that some of the same processes may be at play^102,155–158^. Our findings here suggest that some cell types play an outsized role in landmarks of the midface region and risk for orofacial clefting. Our analysis of marker genes for these specialized subtypes suggests these two subtypes are near one another spatially and could be located near the fusion zone termed the “lambdoid junction”^86,159–161^. Failure of this region to fuse in humans has been suggested to cause cleft clip that could also involve the nostril region and primary palate^10,147,162,163^. It is thus relatively straightforward to imagine that subtle differences in the timing of migrations and fusion of cells residing in this region could influence the shape of the midface. Interestingly, some of the major markers of the specialized ectodermal subtype are multiple members of the EBF family of transcription factors. Our previous work suggested that these genes were co-expressed more strongly in human craniofacial cell types than mouse, and found compelling evidence that *EBF3* is a *bona fide* orofacial clefting risk gene^13^. This EBF family of transcription factors have been linked to regulation of differentiation of multiple different tissue types and predisposition for several tumor types^164–170^. The timing of differentiation of cells at a fusion zone could influence the degree to which structures fuse and impact both clefting risk and facial shape. Studies leveraging the marker genes we have identified for each of these subtypes could allow more specific labelling and identification of these cells in human tissues and mouse embryos as well as experiments to test impact of facial variation.

Our analysis of curated gene lists that are not included in standard gene ontologies was also revealing related to both the cell type identities as well as the composition of the gene lists themselves. For instance, our previous prioritized gene list as well as the curated CleftGeneDB resource are heavily biased toward some mesenchymal subtypes. This is not surprising given the ratios of cell types we observed in the data generated here. Mesenchyme is by far the dominant major cell type, thus previous studies of gene expression and protein expression from bulk tissues were heavily biased toward this cell type. The genes identified as constrained in human populations were more broadly enriched across all the cell subtypes suggesting they play critical roles in most cell types in the body. As expected, the unconstrained genes were enriched in relatively few cell types and were not enriched in the likely craniofacial disease relevant subtypes. While both the common variant analyses for orofacial clefting and the curated craniofacial disease gene lists were biased toward mesenchyme subtypes, genes harboring rare *de novo* protein damaging variants identified in cleft probands showed much more enrichment in ectodermal subtypes. This trend was not observed for *de novo* synonymous variants suggesting this was not a population specific effect or other artifacts of sequencing. This trend was further supported when we examined the frequency of *de novo* protein altering variants, where we found significant enrichment in multiple ectoderm subtypes primarily for CL/P. While the number of CP only cases were fewer than CL/P, we found these *de novo* variants were enriched in a few mesenchymal subtypes that make sense for a spatial perspective. Overall, this points to the ectodermal subtypes, that as we discussed above make up a small proportion of craniofacial tissue, as a major contributor to clefting risk. Due to the biases of previous studies for the most abundant cell types there are likely many additional clefting risk genes that remain to be discovered. The resources we described here could help further prioritize genes that are discovered in such sequencing cohorts. For instance, the nasal placode ectodermal subtype was marked the most substantial number of genes with *de novo* damaging variants. Many known disease risk genes are expressed in this subtype thus genes with similar patterns of expression or specificity of expression could be guilty by association. In particular, our analysis highlighted the *SFRP4* gene. This gene has been linked to Pyle disease (OMIM 265900) that features bone abnormalities and fragility particularly of the long bones and GWAS of bone mineral density^171–174^. Similar phenotypes are observed in *Sfrp4* knockout mice^175^. Cell type specific dysregulation of this gene either due to somatic mosaicism or regulatory element disruption could result in bone abnormalities or other defects in a relevant part of the developing face. Further studies of this gene in a craniofacial specific context in mice as well as identification of the regulatory landscape controlling could reveal a role in clefting risk.

As detailed above, our analysis of craniofacial variation revealed multiple cell types that contribute to human facial shape. Beyond interindividual differences there have been reported to be substantial differences in the shape of many craniofacial features between modern humans and of closely related but extinct hominid species such as Neanderthal and Denisovans^176–178^ . Identifying the genetic contributions to these differences and if Neanderthal derived sequences in the human genome predispose individuals to specific phenotypes or diseases has been of particular interest^179–185^. While Neanderthal derived variants in genes and regulatory regions active in adult bulk tissues have been linked to specific phenotypes related to brain and cranium shape, immunity, and adipose function^186–189^, it is unknown if any human developmental cell types might be influenced by such variants. Our analysis points to Neanderthal derived regions in the European genetic background are systematically enriched near genes with biased expression in multiple cell types related to ear development, cartilage, and specialized ectodermal subtypes. Cartilage1 and EBF expressing ectodermal subtype (ect.EBF) were also shown to be enriched for both *de novo* protein damaging variants in orofacial clefting probands and several aspects of modern human facial variation. These results could suggest that risk for orofacial clefting and facial shape could both be influenced by Neanderthal introgression events. We did not observe any such enrichments for sequences that have been shown to be accelerated on the human lineage, suggesting that findings are functionally relevant. Consistent with this idea, the Neanderthal derived analysis was the only one that demonstrated enrichment in all the ear related ectodermal subtypes. Multiple aspects of Neanderthal inner ear morphology have been shown to differ substantially from modern humans and other primates^190,191^. We also note that we observed enrichment of Neanderthal introgressed regions near genes with biased expression in the specialized ectodermal subcluster ect.GDNF. This was the lone subtype that was enriched for abnormalities of the voice. Differential DNA methylation patterns between modern humans and Neanderthals and Denisovans indicated genes related to vocal anatomy are regulated in a distinct fashion^192^. Thes findings open the distinct possibility that the degree of introgressed segments in the genomes of modern human individuals could influence ear morphology and hearing capabilities as well as vocal characteristics.

In summary we have provided a substantial resource for understanding the cell types and gene expression patterns that build the human and mouse face. Our analyses revealed relationships between specific cell subtypes and many aspects of human biology including facial shape and orofacial clefting risk. We also illuminated potential contributions of ancient hominids to craniofacial morphology. Future integration with cell type specific chromatin accessibility could reveal specific variants and regulatory regions that encode such phenotypic differences, risk factors, and species-specific biology. This data can be explored through an interactive web application that is accessible to most researchers: https://cotneyshiny.research.chop.edu/shiny-apps/craniofacial_all_snRNA/. The data will be deposited to other major single cell aggregation databases including the Chan-Zuckerberg CellXGene Discover resource ^39,193^.

## Acknowledgements

We would like to thank members of the UConn/JAXGM Single Cell Genomics Core for help with standardizing single-cell isolation techniques and preparing sequencing libraries. We would also like to thank members of the UConn Computational Biology Core and High-Performance Computing Facility for assistance with package installation and software/hardware support. We are grateful to Dr. Peter Tran, Dr. Sungryong Oh, and Pooja Sonawane for constructive feedback and copyediting. This work was funded by grants from the National Institutes of Health to E.J.L.C (R01-DE030342, X01-HG010835, X01-HD100701, X01-HL132363) and JC (NIDCR 1R01DE028945, NIDCR 1R03DE028588, and NIGMS 5R35GM119465).

## Author contributions

Conceptualization: J.C. Investigation: N.F., E.W.W. and J.C. Formal analysis: N.F., E.W.W, B.M.S. K.R., S.W.C, J.C. Writing—original draft: J.C. Writing— review and editing: N.F., E.W.W, B.S, K.R., S.W.C, N.G.S, E.J.L.C., J.C. Funding acquisition: J.C. Supervision: J.C., E.J.L.C., S.G.K.

## Code and data availability

Code for analysis and generation of figures can be found on github (https://github.com/cotneylab/craniofacial_snrna). An interactive website for exploring processed data is found here: https://cotneyshiny.research.chop.edu/shiny-apps/craniofacial_all_snRNA/. Raw data from mouse experiments generated in this work will be deposited in GEO. Cellranger ARC gene expression outputs for both human and mouse are available on Zenodo.

## Methods

### Human tissue samples

The use of human embryonic tissue was reviewed and approved by the Human Subjects Protection Program at UConn Health (UCHC 710-2-13-14-03) and Children’s Hospital of Philadelphia (IRB 24-022258). Human embryonic craniofacial tissues were collected via the Joint MRC/Wellcome Trust Huma Developmental Biology Resource (HDBR) under-informed ethical consent with Research Tissue Bank ethical approval (18/LO/0822 and 18/NE/0290, project 200225). Donations of tissue to HDBR are made entirely voluntarily by women undergoing termination of pregnancy. Donors are asked to give explicit written consent for the fetal material to be collected, and only after they have been counseled about the termination of their pregnancy. Further documentation of all policies and ethical approvals for HDBR sample collection can be found at https://www.hdbr.org/ethical-approvals. Tissues were flash-frozen upo collection and stored at −80 °C. Upon thawing, the samples were quickly inspected for intactness of the general craniofacial prominences and processed for single nucleus multiomics.

### Mouse embryonic tissue samples

The use of mouse embryonic tissues was reviewed and approved by the UConn Health Institutional Anim Care and Use Committee (Protocol AP-2000061-0723). Eight-week-old wild-type male and female C57BL6/J mice were obtained from Jackson Laboratory. Mice were housed according to recommendatio by Jackson Laboratory with 12 h light:dark cycle beginning at 7 a.m. The ambient temperature was maintained between 20 and 22 °C and humidity was maintained at 40–60%. Mice were given ad libitum access to food and water. Timed matings were established by the identification of vaginal plugs the morning following the housing of a single male with multiple female mice. Embryos were harvested from pregnant mothers at mid-day either 10, 11, or 12 days after identification of the vaginal plug. The staging was confirmed by counting somites and comparing overall morphology to the Theiler Staging Criteria^194^. embryos from a given litter were combined for individual biological replicates, and at least three biologica replicates were collected and processed for each stage. Craniofacial prominences were collected in a ve similar fashion to human samples and subsequently prepared for single nucleus multiomics.

### Single nucleus multiomics

Primary human craniofacial tissues from CS12, CS13, CS14, CS16, CS17 and CS20, each stage represented by a minimum of 3 replicates, were obtained from HDBR. Tissue from each embryo were mechanically broken into single-cell suspensions and cells were checked for viability counted using Tryp blue staining following the 10X Genomics protocol for single-cell multiome sequencing using the ChromiumX controller. Samples were sequenced on multiple Illumina NovaSeq runs according to 10X Genomics recommendations. Raw fastqs were processed using CellRanger ARC (v2.0.2) using hg38 genome and gene annotations provided by 10X Genomics.

Primary mouse craniofacial tissues from E10.5-E12.5 from multiple (3-5 depending on stage) mixed sex C57BL/6J Mus Musculus embryos (Jackson Laboratories) were pooled. Animals were raised and sacrificed in compliance with UConn Health IACUC approval (protocol AP-200061-0723). Samples were mechanically broken into single-cell suspensions, processed for multiome using the ChromiumX controll and sequenced in the same fashion as for human samples above. Raw fastqs were processed using CellRanger ARC (v2.0.2) using mm10 genome and gene annotations provided by 10X Genomics.

### Processing of snRNA and identification of major cell types

Filtered barcode matrices from each human samples generated by CellRanger ARC were individually loaded with Read10X_h5 command in Seurat^195^ and merged into one object. Percentage of mitochondri reads were calculated for each cell and filtering was performed to only retain cells with less than ten percent mitochondrial derived. Further filtering was performed based on number of counts per cell (500 < < 25000) and number of genes detected per cell (500 < x < 7000). Filtered data were normalized with default values and cell cycle scores were calculated using Seurat. Data was scaled based on S and G2M score regression and dimensionality reduction with principal component analysis (PCA) were performed using respective commands in Seurat. The top 2000 variable features were identified and data were the further integrated with harmony R package^196^. Nearest neighbors based on harmony corrected embeddings were calculated with up to 30 dimensions and clusters were identified with multiple resolutio from 0.1 to 1 in Seurat. We then performed uniform manifold approximation and projection (UMAP) dimensionality reduction using harmony corrected embeddings in Seurat (dimensions = 30, minimum distance = 0.3). Resulting clusters were inspected for expression of multiple craniofacial markers form Li al 2019 and marker genes were identified for each cluster. Cells from clusters identified with high expression of neuronal markers *TUBB3* and *MAP2* were removed and the process of normalization, harmonization, and clustering was repeated with remaining cells from all samples. Marker genes for maj cell types were identified using FindAllMarkers (logfc.threshold = 0.25, min.pct = 0.1, test.use = “wilcox”, min.cells.feature = 3, mi.cells.group = 3, pseudocount.use =1, and return.thresh = 0.01). The top 100 marker genes for each cluster (p < 0.05, ranked by log2fold change versus all other clusters) were then analyzed for gene and disease ontology enrichments using compareCluster in clusterProfiler R package 4.12.6). Cranial neural crest genes were compiled based on markers identified by regulatory network construction in human cultured CNCC and craniofacial tissue data^69^. Major cell type labels were applied each cluster.

For mouse data sets, filtered barcode matrices from each mouse E10.5 to 12.5 samples generated by CellRanger ARC were individually loaded with Read10X_h5 command in Seurat^195^ and merged with E13 to E15.5 data from Pina et al 2023 (GSE205448). Calculation of percent mitochondrial reads and filtering were performed with similar thresholds to human data above. Subsequent harmonization, dimensionality reduction, and clustering were performed identically to those for human data above. Less significant contamination of neuronal cell types was observed in mouse data, which was identified and filtered as describe for human. Identification of marker genes and gene ontology enrichments, and CNCC modules scores were performed as above for human data. Major cell type labels were applied to each mouse cluster.

### Marker gene comparisons across species

Lists of all marker genes for each of the seven main subtypes for each species (p < 0.05) were compiled and orthology based on HGNC symbol annotated by Ensembl v105 was obtained using the getLDS command in the biomaRt R package (v. 2.60.1). Only genes that had one ortholog in each species and a HGNC symbol were retained (n = 7504). Significant overlaps between all orthologous marker gene lists were determined using the testGeneOverlap command in GeneOverlap R package^197^ Conserved and species-specific genes were determined based on HGNC symbol and the intersection matrix obtained by getMatrix in GeneOverlap. Gene and disease ontology enrichments were calculated using clusterProfiler Final Seurat objects were prepared for display in an interactive webapp using the ShinyCell R Package^19^

### Subclustering of major cell types

For major cell types labelled as mesenchymal, ectodermal, or CNCC further subclustering was first performed on mouse data. For each major cell type, normalization, scaling with regressed cell cycle impacts, harmonization, and subclusters were identified using same procedure as described above. Marker genes were identified for each cluster and functional enrichments were determined using clusterProfiler. Annotations for each cluster were manually assigned based on those originally described^16^. Mouse cell subtype assignments were further confirmed with ToppGene^199^ using the scToppR package^200^. Mouse main and subtype annotations were further confirmed by projection on mouse E15.5 spatial transcriptomics data^23^ (GSE245469). Links between snRNA and spatial data were determined using FindTransferAnchors and transferred using TransferData in Seurat.

Following annotation of subtypes and for comparison with human data, an intermediate data set was created where mouse genes were reduced and converted to those to those with one to one orthology wit human genes using annotations provided by Ensembl (archive dec2021) with biomaRt R package^201^. Th intermediate dataset was used to transfer mouse subtype annotations to human subtypes by first identifying shared features across clusters using FindTransferAnchors in Seurat with log normalization a canonical correlation analysis (cca). Predicted subtype labels were transferred to human subtypes using TransferData in Seurat and further confirmed with a confusion matrix. Marker genes for human subtypes were identified as performed for major cell types and functional enrichments were characterized with compareCluster in clusterProfiler. Final Seurat main objects and subtype objects were prepared for displ in an interactive webapp using the ShinyCell R Package^198^. Seurat objects were also converted to scanp and anndata objects using scEasy R package (v0.0.7) for hosting at the Chan Zuckerberg CELL by GEN Discover resource.

### Processing of spatial transcriptomics

Spatial transcriptomics data for two sections of a human CS13 embryo^25^ were retrieved from https://heoa.shinyapps.io/code/. Raw sequence count matrices were loaded using the Read10x command of Seurat^195^ and converted to HDF5 format. These counts were then combined with spot coordinates and section images using CreateSeuratObject. Data from both slices were merged and variable features wer determined using Seurat. The percentage of mitochondrial reads was determined for each cell and was used to transform all data in the merged object using SCTransform from Seurat. Data was clustered usin UMAP and plotted which revealed a strong batch effect between the two spatial objects. Data was furthe normalized using Harmony (v1.2.3), projected using UMAP, and clustered with a resolution of 0.8. Marke genes for the 22 clusters were identified using FindAllMarkers in Seurat. The top 100 marker genes for each cluster (p < 0.05, ranked by log2fold change versus all other clusters) were then analyzed for gene and disease ontology enrichments using compareCluster in clusterProfiler R package (v. 4.12.6). Enrichments and spatial localization were compared to previous annotations by Xu et al 2023 and labelle accordingly. The top 100 marker genes from each of the subclusters identified in human craniofacial dat were used to calculate module scores across the merged spatial object and plotted using SpatialFeaturePlot in Seurat.

We chose mouse E11.5 spatial transcriptomics data as it is most morphologically similar to CS13 human embryos. Data were retrieved all E11.5 spatial transcriptomics data from the MOSTA resource^43^ and loaded into Seurat as for human data above. For each section, module scores of top 100 marker genes f each main cluster or subcluster were calculated. Gene spatial feature plots for selected genes and modu scores were then generated with Seurat.

### Facial variation and congenital abnormality GWAS enrichments

We retrieved summary statistics for facial variation^87,88^ and all congenital abnormality GWAS summary statistics from FinnGenn^116^. Raw summary stats were further processed and standardized with hg38 cooridinates with MungeSumstats R package (https://doi.org/10.1093/bioinformatics/btab665). Variants were mapped to genes +/-100kb using MAGMA^89^Frontal facial measures and FinnGenn were processe based on 1000 genome European population while profile facial measures were processed with the 100 genome Middle/South American population all obtained from the MAGMA website (https://cncr.nl/research/magma/). We converted the Seurat snRNA-seq expression data to a CellTypeData set with the Expression Weighted Celltype Enrichment (EWCE) R package^90^ and then assessed each study trait for a linear positive correlation of cell type gene expression specificity and gen level genetic associations using MAGMA Celltyping^91^. Plots were generated using tidyheatmaps in R^202^.

### Gene list enrichments per cell type

The CellTypeData-formatted human craniofacial snRNA-seq objects were generated using generate_celltype_data in EWCE (v1.15.0). Mean and specificity metrics for several marker genes (*SOX TP63*, and *MSX1*) were inspected across main cell types and subtypes using plot_ctd in EWCE. Gene lis were compiled from multiple resources including gnomAD (v4.1), CleftGeneDB, prioritized genes and bla module from craniofacial WGCNA^13^, and genes affected *by de novo* variation in orofacial clefting probands^32,123^. For Neanderthal introgressed regions and human accelerated regions, coordinates were obtained from respective publications^188,203^ and assigned single nearest gene using rGREAT^204^ with “oneClosest” association rule. Each gene list was then tested for linear association using bootstrap enrichment test in EWCE (reps = 10,000; geneSizeControl = TRUE). Results from all gene lists were the merged and plotted with ewce_plot in EWCE with correction for total number of gene lists and cell types tested using the Benjamini-Hochberg approach.

### Phenotype-cell type association tests

To map the relationships between cell types and phenotypes, we ran pairwise association tests between all combinations of cell types in our snRNA-seq-derived CellTypeData and phenotypes across the Huma Phenotype Ontology (HPO)^119^ using the run_phenomix function from MSTExplorer (v1.0.5). In contrast t the gene list-based approaches (e.g. EWCE) this function reframes the problem as a series of linear regressions by leveraging continuous scores that summarize the current strength of evidence for a caus relationship between each gene-phenotype pair (using additional data from the Gene Curation Coalition)^118,205^. The continuous nature of this data allows us to more accurately capture phenotype-cell type relationships, especially for phenotypes with large gene lists where only some genes have strong evidence of actually causing the phenotype. The gene signature vectors for each phenotype were previously merged and shared as a single precomputed gene (5003 unique gene symbols) x phenotype (11047 unique HPO phenotypes) association matrix. Next, a series of linear regressions tests were performed between the gene specificity vectors of each cell type (n=66 vectors) and the gene associatio vectors of each phenotype (n=11047 vectors). Finally, multiple-testing correction was applied using Benjamini-Hochberg False Discovery Rate^206^ (at FDR<5% significance).

For the purposes of summarization and visualization, the number of significantly associated phenotypes per cell type were then computed within each major HPO branch (Fig. 7C). Here, we define HPO branch as groups of related phenotypes that can be labeled according to their shared ancestral term, e.g., ‘Abnormality of the immune system’. Next, we sought to determine whether some cell types were disproportionately more often associated with phenotypes of a particular HPO branch. To accomplish thi we performed a series of proportion tests comparing the proportion of total phenotypes that a given cell type was significantly associated with within a target HPO branches relative to all other HPO branches. I practice, we computed 2×2 contingency tables (number of significant phenotype association vs. number non-significant phenotype associations x target branch vs. non-target branches) for each cell type within each HPO branch, which were then used as inputs to the prop_test function within the rstatix R package (v0.7.2). This test appropriately takes into account the different number of phenotypes across HPO branches. Only one-sided tests were performed to test whether the target HPO branch was greater than other (non-target) branches (set with the alternative = “greater” parameter). All proportion tests were the corrected for multiple testing at FDR<5%.

### Orofacial Clefting de novo variant analysis

We used the R package ‘DenovolyzeR’ (version 0.2.0) to test enrichment of *de novo* variants (DNs) in a dataset of OFC case-parent trios. Enrichment is calculated by comparing the expected number of variants, as determined by mutation models described by Samocha, et al^130^, to the observed number of variants in a given gene or group of genes using the ‘DenovolyzeByClass’ and ‘includeGenes’ functions. Using our dataset of 2031 DNs in 1171 genes identified in 1676 trios with OFCs, we first compared this list of genes to those with calculated mutational rates in the R package ‘DenovolyzeR’ (version 0.2.0) using the ‘viewProbabilityTable()’ function. There were 12 trios in which DNs were identified, but no mutational rates for the affected genes were present; thus, we ultimately tested 1662 trios with OFCs, broken down by subtype including 1180 cleft lip with or without cleft palate (CL/P; 226 cleft lip (CL), 954 cleft lip and palate (CLP)), and 482 cleft palate (CP) trios. We then tested enrichment of all OFC trios and by subtype within the top 20% of genes by log2FC derived from single nucleus RNA sequencing of human craniofacial tissue at CS20.

